# Cross-comparison of state of the art neuromorphological simulators on modern CPUs and GPUs using the Brain Scaffold Builder

**DOI:** 10.1101/2022.03.02.482285

**Authors:** R. De Schepper, N. Abi Akar, T. Hater, B. F. B. Huisman, E. D’Angelo, A. Morrison, C. Casellato

## Abstract

A variety of software simulators exist for neuronal networks, and a subset of these tools allow the scientist to model neurons in high morphological detail. The scalability of such simulation tools over a wide range in neuronal networks sizes and cell complexities is predominantly limited by effective allocation of components of such simulations over computational nodes, and the overhead in communication between them. In order to have more scalable simulation software, it is therefore important to develop a robust benchmarking strategy that allows insight into specific computational bottlenecks for models of realistic size and complexity. In this study, we demonstrate the use of the Brain Scaffold Builder (BSB; De Schepper et al., 2021) as a framework for performing such benchmarks. We perform a comparison between the well-known neuromorphological simulator NEURON (Carnevale and Hines, 2006), and Arbor (Abi Akar et al., 2019), a new simulation library developed within the framework of the Human Brain Project. The BSB can construct identical neuromorphological and network setups of highly spatially and biophysically detailed networks for each simulator. This ensures good coverage of feature support in each simulator, and realistic workloads. After validating the outputs of the BSB generated models, we execute the simulations on a variety of hardware configurations consisting of two types of nodes (GPU and CPU). We investigate performance of two different network models, one suited for a single machine, and one for distributed simulation. We investigate performance across different mechanisms, mechanism classes, mechanism combinations, and cell types. Our benchmarks show that, depending on the distribution scheme deployed by Arbor, a speed-up with respect to NEURON of between 60 and 400 can be achieved. Additionally Arbor can be up to two orders of magnitude more energy efficient.

## 1 INTRODUCTION

The role of computational methods in the neuroscience is well established, with various large funding frameworks (Human Brain Project, BRAIN Initiative, Brain/MINDS, China Brain Project, Blue Brain Project, etc.) allocating significant fractions of their budgets towards the accurate simulation of neuronal signalling in brain and nerve tissues (Einevoll et al., 2019). The computational complexity of these neuronal networks remains a challenge: simulating a whole human brain would, according to some calculations, require up to ∼ 4 × 10^29^ TFLOPS (Fan and Markram, 2019). For comparison, the first place in the November 2021 TOP500 ranking of the world’s supercomputers is held by the Fugaku Supercomputer at RIKEN, Japan, with a peak performance of 5.4 × 10^5^ TFLOPS. Even if the level of detail in the simulation can be significantly reduced, the full simulation of the human brain will remain out of reach for a good while to come. The largest simulations of spiking neural networks to date are ∼ 1.86 × 10^9^ nodes, connected by ∼ 11.1 × 10^12^ synapses for a network of point neurons (Kunkel et al., 2014), and ∼ 1.0 × 10^7^ nodes for a network with morphologically detailed cells (Reimann et al., 2019). This gap in required and available compute power places significant limitations on the possibilities of computational neuroscience research, and improvements in implementation of simulation codes help address this problem.

The complexity of a complete neuroscientific model is to a large extent determined by the detail of the biophysical processes present in the model of the cell and the size and detail of the morphology, which prescribes the number of compartments to simulate. Together with the size of neuronal network (i.e. the number of neurons) these factors determine the required computational power for simulation. The impact of the size of the network is relatively well understood and largely scalable with available computational resources (and time), but the impact of the complexity of biophysical mechanisms has not been studied to the same degree.

To address this issue, we extended the Brain Scaffold Builder (BSB) to be able to export simulations to Arbor, in addition to the existing export to NEURON. This effort is motivated by preliminary experiments showing significant performance improvements using Arbor. We benchmark 1. a collection of biophysical mechanisms, 2. a collection of cerebellar single cell models, and 3. two differently sized networks representing a cerebellar cortex model on various multi-node and multi-GPU hardware configurations. Studying performance gains at these levels of a neuroscientific simulation enables us to deconstruct the performance of a full model.

Compared to NEURON, a speed-up of at worst a factor two and at best factor 400 support the thesis that Arbor delivers on the promise of increased computational efficiency. Reducing the time to solution saves on scientific resources, which means more and more thorough simulation experiments can be performed for the same compute time budget available to a researcher. Moreover, it allows previously unfeasible experiments to be conducted, and thus opens the door to new research questions.

### 1.1 State of the Art

The NEURON simulator was initially established in 1984 (Hines, 1984). Whereas alternatives such as Genesis (Bower et al., 2003) and Moose (Ray et al., 2008) exist, they have not generated the same level of developer and user engagement, leading to NEURON’s status as the gold standard simulator for networks of multi-compartmental neurons. Various versions of NEURON have forked in order to address the challenges of building and running large networks on High Performance Computing (HPC) and new accelerators; CoreNEURON (Kumbhar et al., 2019) and NeuroGPU (Ben-Shalom et al., 2020) perhaps being the most well known of these offshoots. CoreNEURON shows it reduces memory consumption by a factor of 4-7, and execution times are reduced 2 to 7-fold. In addition to adaptation to HPC and new hardware environments, there has been increased interest in developing an alternative technology path to NEURON, leading to new multi-compartmental simulators such as EDEN (Panagiotou et al., 2021) and Arbor (Abi Akar et al., 2019). These break with NEURON compatibility in part or full, not only to enable better computational efficiency, but also to favor specification based formats like NeuroML (Cannon et al., 2014), which are easier to write and comprehend, and, because of their formal specifications, enable better interoperation with the neuroscientific software landscape. Moreover, formal specifications play an important role in making it feasible to write new simulator implementations from scratch.

The Brain Scaffold Builder is the first tool to support complex networks in both Arbor and NEURON, making direct comparisons possible. The BSB is a declarative framework aimed at helping neuroscientists write organised explicit code, with all aspects of the network description visible at a glance in a human readable configuration file. It excels at biophysically detailed models but the paradigms have been applied to mesoscale and macroscale projects as well. It offers out of the box solutions for common modelling problems such as N-point problems of cells in space, morphologies, placement and connectivity problems. In summary, it aims to simplify setting up correct many-component neuronal network reconstructions and simulations.

## 2 METHODS

Comparisons across simulators of the same biologically realistic models have a few advantages over traditional benchmarks of idealized situations such as ring networks. Biologically realistic models better represent the workload, require and rely on a larger collection of features, and cover more code of the simulator, thus better stress testing the stability and quality. In this publication we will use the BSB to generate such biologically realistic models for two simulators, NEURON and Arbor, and so assess their respective performances. We compare all the components and subsystems making up the multi-compartmental network model of the cerebellar cortex individually.

First we consider the membrane mechanisms, the elemental building blocks that determine the membrane potential fluctuations of each compartment of a single cell. Next, multiple compartments with multiple mechanisms, as prescribed by the cell morphology and electrophysiology, are linked together to form a single cell. Finally, all these single cells are put in a network together, by communicating with another through synapses.

To create equivalent network models in NEURON and Arbor, we added Arbor support to the BSB. Support was included for input devices (such as spike generators), morphologically detailed cells, connections, gap junctions, output devices (called *probes* in Arbor); and the full BSB simulation interface was implemented allowing the framework to run fully managed Arbor simulations on a single thread, multiple threads, GPU backends or distributed in HPC environments commonly using Message Passing Interface (MPI).

In order to obtain a more fine-grained analysis of the performance of the simulators on separate tasks of the simulation, we further subdivided the network stage into the largest network that could be supported by a single unit of hardware (workstation or HPC compute node) and a full scale benchmark distributed over separated units of hardware. A detailed technical description to set up the benchmarks can be found in the Data Availability Statement.

### 2.1 Mechanisms

The cell membrane and the many kinds of biophysical mechanisms present thereon (e.g. sodium ion channels) are responsible for the propagation of action potentials in nervous tissue and as such thought to be the substrate of the computing power of nervous and brain tissue. The cell membrane can be modelled as a set of mathematical differential equations, each representing a difference process in the phospholipid bilayer. Solving the system of equations allows one to evolve the state of the system: to propagate action potentials.

We adapted a catalogue of 33 NEURON NMODL mechanisms^1^ to the Arbor dialect^2^. The mechanisms^3^ were collected from the granule, Golgi, Purkinje, basket and stellate cell models^4^ used in the model of the cerebellar cortex (Masoli et al., 2020, 2021; Rizza et al., 2021; Masoli and D’Angelo, 2017).

We carried out the mechanism benchmarks on a single thread on an AMD Ryzen 7 3700X 8-Core Processor. One of 46, 939 combinations of zero to four mechanisms were inserted on a single compartment and repeated 30 times each. Each combination was simulated for 10 ms with an integration timestep of 0.025 ms. Some combinations of mechanisms are invalid, for example when they write information to the same ion species. Combinations of mechanisms that interact with the same ion species are excluded from the tests.

All kernel density estimations (KDE) were estimated using scipy.stats.gaussian kde, which was provided solely with the samples, relying on Scott’s rule for bandwidth estimation (Scott, 2015).

### 2.2 Single cell models

Single cell benchmarks were also run on a single thread on an AMD Ryzen 7 3700X 8-Core Processor, 3593 MHz, integrating the mechanisms into a functional unit with validated electrophysiology (Masoli et al., 2020, 2021; Rizza et al., 2021; Masoli and D’Angelo, 2017)^5^. The Arborize^6^ package was used to write cell model descriptions in a single format that can construct equivalent cells in both Arbor and NEURON. The performances of the cell models were compared by running each model 30 times for 1000ms with an integration timestep of 0.025 ms.

### 2.3 Network model

We performed benchmarks for two sizes of cerebellar cortex model: a small model with 1, 000 cells and 10, 000 synapses, and a large model with 30, 000 cells and 1.5 million synapses (De Schepper et al., 2021). Each type of benchmark was repeated ten times for 1000 ms with an integration timestep of 0.025 ms. The average firing rates of the populations were compared and raster plots are provided in the Supplementary Material.

As published, the cerebellar cortex model uses complex synapses (AMPA, NMDA, and GABA) which implement explicit spike history buffers to approximate solutions to differential equations (Nieus et al., 2005). This method does not scale to large simulations, because of the increase in computational complexity and memory requirements. A better approach, based on direct integration of the ODEs, is in development. For this discussion, these complex synapse types have been replaced with simpler exponential synapses in the model.

#### 2.3.1 Network benchmarks

As performance metrics we consider a) the time-to-solution, i.e. the wall-clock time of a simulation, and b) the timestep duration, i.e. wall-clock seconds it takes to solve a biological millisecond (s_wall_*/*ms_bio_). Simulations were executed on PizDaint^7^ supercomputer. Timestep duration is in current morphologically detailed simulators orders of magnitude more than the biological time simulated, presenting us with an obvious shortcoming of such simulations. A final consideration is the total energy cost of the simulation, as ecological concerns grow. PizDaint’s workload manager, Slurm, provides an energy consumption estimate for each workload, which is used to determine the total energy cost of a simulation. Each simulation is executed ten times, ensuring a representative mean and provides an estimate for the standard deviation.

NEURON does not support multi-threaded network simulations or GPU accelerated simulations, but does support MPI distribution. We carry out single-threaded, non-distributed simulations of the small 1, 000 cell model in NEURON and compared to single-threaded, multi-threaded, hyper-threaded, and GPU-accelerated equivalent simulations in Arbor. Distributed MPI simulations of the regular 30, 000 cell model are carried out in NEURON, on 20 nodes with 36 MPI ranks each, resulting in a distribution of the model over 720 MPI processes and a cost of 20 node hours per wall-clock hour (*NEURON*). The results are compared to equivalent 720 process MPI-distributed simulations (*Arb. MPI*, multi-threaded simulations on 20 nodes with 2 MPI ranks per node and 18 threads per MPI process (*Arb. multithr*.), and hyper-threaded simulation with 36 threads per MPI rank (*Arb. hyperthr*.). The NEURON results are further compared to GPU-accelerated simulations on one to 20 GPU nodes, with 1 Nvidia Tesla P100 GPU per node (*Arb. GPU*).

For the distribution across multiple nodes, it’s important to consider the underlying hardware architecture. PizDaint features CPU compute nodes with two sockets of 18 physical cores (and 36 logical cores) each. Arbor’s optimal configuration for a single socket is multithreading, and to use MPI between sockets, local or remote. NEURON does not have a multithreaded mode, so we compare to a fully MPI-distributed NEURON simulation. Arbor can be configured similarly (MPI on socket instead of multithreading, labelled *Arb. MPI*), which is added to the comparison matrix to ensure the fairest and most informative comparison.

The network models were constructed using the methods described in De Schepper et al. (2021), resizing the network description to obtain suitable estimated computational loads for the benchmarks in terms of cell and synapse number. A BSB-to-Arbor adapter was developed which can create Arbor’s functional *recipes*. When Arbor calls these recipe functions, the adapter looks up the required information in the BSB network description. By invoking the BSB’s NEURON and Arbor adapters the same network reconstruction could be instantiated in both NEURON and Arbor for exact comparisons.

## 3 RESULTS

The mechanism catalogue benchmarks yielded speed-up factors for Arbor between 3 to 20 times faster than NEURON (Fig. 1 left) with outliers (not shown) up to 30 times faster. Investigation of the multi-modal distribution (see Suppl. Fig. S1) showed ten mechanisms whose speedup distribution lacked a tail with results above 10 times speedup, which the other mechanisms do have. The distribution of maximum speedup factors (Fig. 1 center) for each member in a group shows that benchmarks where elements with non-linear dynamics or a high number of RANGE variable declarations in their NMODL occur are bound to a maximum speedup of less than ten, while benchmarks containing regular elements can be 10 to 25 times faster^8^.

**Figure 1.**
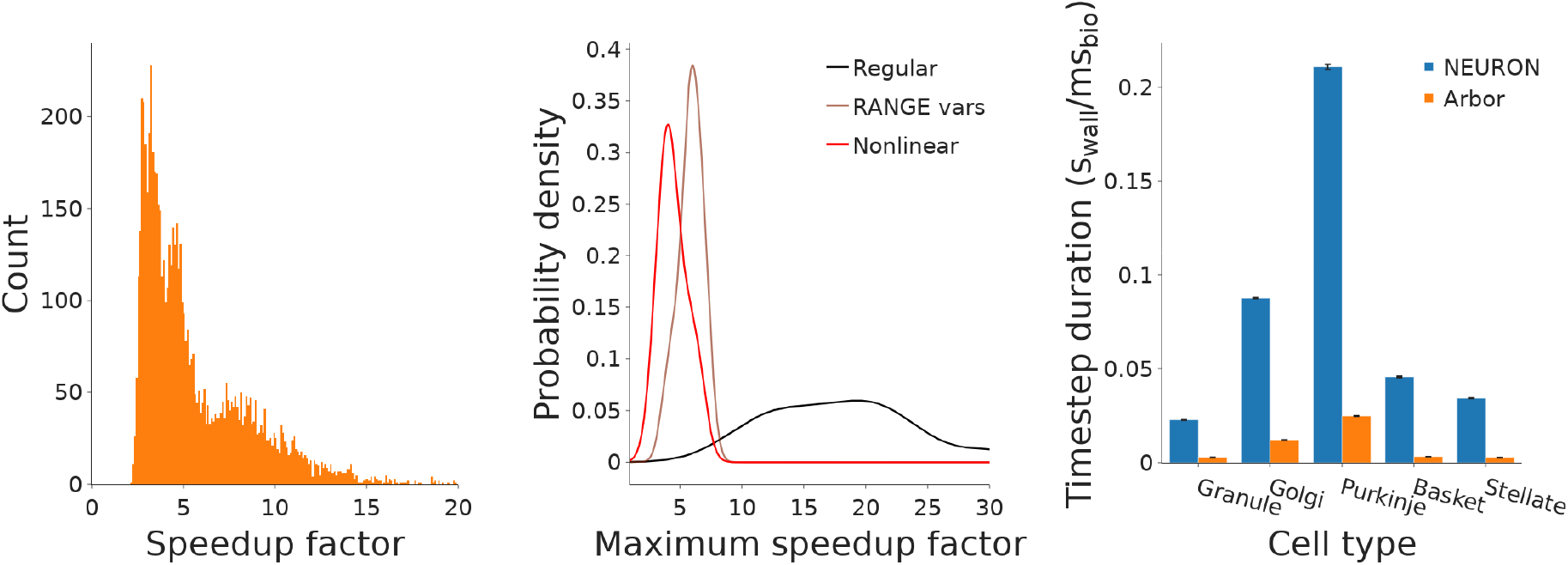
Left: Histogram of the speed-up factors for Arbor with respect to NEURON of 46, 939 mechanism combinations inserted in a single compartment. Center: Kernel density estimates of the maximum speedup of 3 groups of mechanisms: mechanisms with nonlinear dynamics, such as cdp5, mechanisms with high amounts of RANGE variables, and all other mechanisms lacking extraordinary NMODL features. Right: Comparison of the timestep duration for NEURON and Arbor to solve biological timesteps of the cerebellar single cell models.

The single cell models show a speed-up factor of 7 to 12 for Arbor with respect to NEURON (fig. 1 right). The size, mechanism makeup, and estimated computational load of the models can be seen in Suppl. Table S1. The obtained speedups between the simulators are all in the same range despite variations in the cell model sizes and used mechanisms.

The benchmark metrics in table 1 (section: *Undistributed*) of the undistributed scaled-down network model of 1, 000 cells (Fig. 2) show that Arbor on a single thread, refraining from using any technique that NEURON does not support, scores ten times better on all metrics than NEURON, and this gain can be improved to two orders of magnitude using multiple threads. Using GPU acceleration, Arbor advances biological times 400 times faster and uses a hundred times less energy than NEURON. Multiplying the timestep duration with the biological duration of the simulation gives the total time spent simulating; doing this for the undistributed cases yields 100, 57, 65, and 54 seconds spent outside simulation, for NEURON, Arb. single, Arb, multithr. and Arb. hyperthr., respectively. For the NEURON and single-threaded case this is 1% and 8% of the total time, but for the multi-threaded and GPU-accelerated cases this is the larger part of the total task and shows the importance of the way the simulator interfaces with the user, and to promote and enable performant coding. In absolute terms the BSB was able to set up this model twice as fast in Arbor than in NEURON.

**Table 1.** Overview of the measured benchmark metrics. Highlighting indicates the best performance in each section.

| | Time-to-solution (s) | Timestep duration ($s_{\text{wall}}/\text{ms}_{\text{bio}}$) | Energy (kJ) |
| --- | --- | --- | --- |
| **Undistributed** |  |  |  |
| NEURON | 65000 ± 4000 | 64 ± 4 | 750 ± 50 |
| Arbor Singlethreaded | 7120 ± 60 | 6.55 ± 0.06 | 82 ± 2 |
| Arbor Multithreaded | 830 ± 40 | 0.248 ± 0.003 | 12.1 ± 0.3 |
| Arbor Hyperthreaded | 830 ± 50 | 0.1828 ± 0.0002 | 11 ± 0.5 |
| Arbor GPU | 700 ± 30 | 0.166 ± 0.001 | 6 ± 0.2 |
| **Single Socket** |  |  |  |
| NEURON | 5200 ± 200 | 3.6 ± 0.2 | 94 ± 4 |
| Arbor | 1070 ± 30 | 0.4706 ± 0.0006 | 13.8 ± 0.4 |
| **Distributed** |  |  |  |
| NEURON | 17400 ± 300 | 16.7 ± 0.2 | 93000 ± 2000 |
| Arbor MPI | 3600 ± 600 | 2.68 ± 0.05 | 18000 ± 1000 |
| Arbor MPI & multithr. | 2270 ± 20 | 1.99 ± 0.02 | 6500 ± 100 |
| Arbor MPI & hyperthr. | 2700 ± 200 | 2.16 ± 0.03 | 7500 ± 400 |
| **Arbor GPU** |  |  |  |
| 1 node | 4090 ± 30 | 3.56 ± 0.02 | 625 ± 14 |
| 2 nodes | 2320 ± 10 | 1.78 ± 0.01 | 617 ± 8 |
| 4 nodes | 1245 ± 3 | 0.902 ± 0.003 | 632 ± 7 |
| 8 nodes | 760 ± 20 | 0.3855 ± 0.0003 | 680 ± 10 |
| 12 nodes | 590 ± 20 | 0.2773 ± 0.0004 | 790 ± 20 |
| 16 nodes | 650 ± 50 | 0.276 ± 0.001 | 1090 ± 50 |
| 20 nodes | 720 ± 80 | 0.2759 ± 0.0007 | 1420 ± 110 |

**Figure 2.**
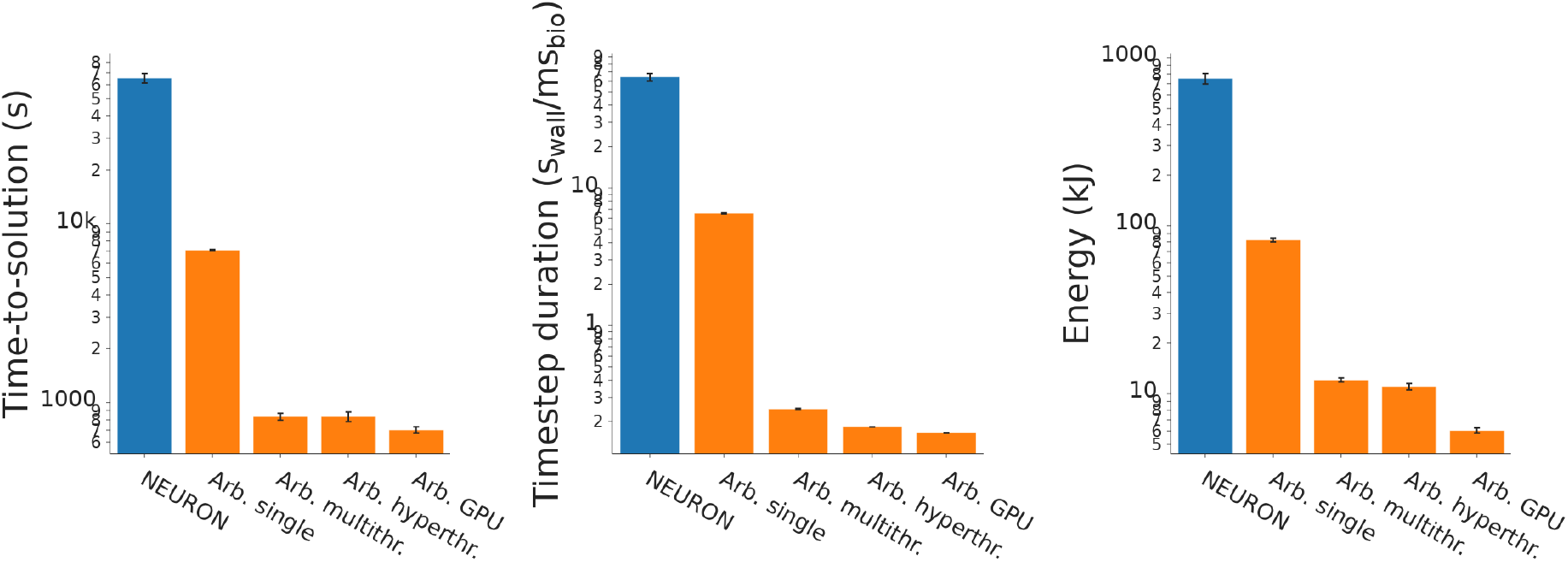
Single machine network model simulations. Time-to-solution (left), timestep duration (center), and energy consumption (right). *NEURON* and *Arb. single* bars show the single threaded benchmarks, *Arb. multithr*. and *Arb. hyperthr*. the multi-threaded and hyper-threaded benchmarks, and *Arb. GPU* the GPU-accelerated benchmarks.

Using Arbor’s optimal hardware configuration on PizDaint (multithreading on a socket, MPI between sockets/nodes) show that Arbor’s capacity to utilize multicore architectures lead to a performance improvement over NEURON of a factor of five to ten for all metrics (Fig. 3 and table 1, *Single Socket*).

**Figure 3.**
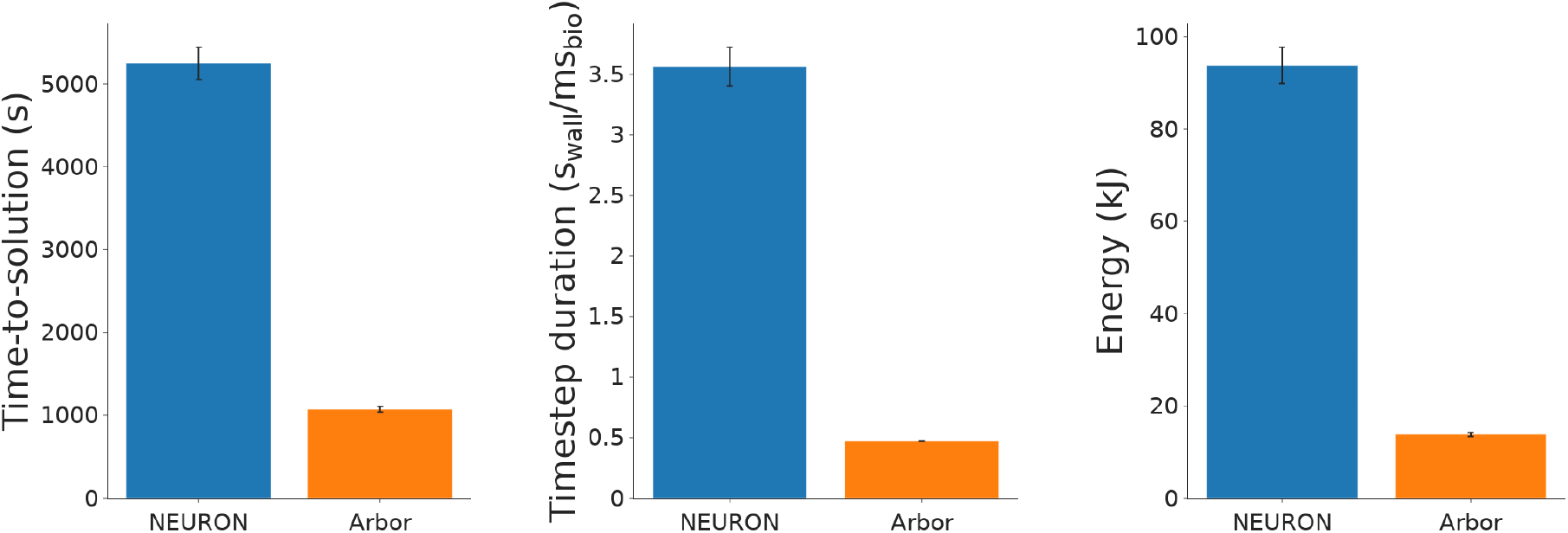
Comparisons of the time-to-solution (left), timestep duration (center), and energy consumption (right) of the single socket network model simulations.

The full scale 30, 000 cell network was distributed over 20 CPU nodes (Fig. 4 and table 1, *Distributed*). Arbor exhibited a speed-up with respect to NEURON of about five times across all metrics when restricted to parallelize over MPI (*Arb. MPI* bars). When parallelization schemes better suited to the HPC architecture are used, the performance across all metrics doubles to eight times better time-to-solution and timestep durations for Arbor, and is also 14 times more energy efficient (Fig. 4, *Arb. multithr*. bars). Enabling hyperthreading did not lead to any further performance improvements (Fig. 4, *Arb. hyperthr*. bars).

**Figure 4.**
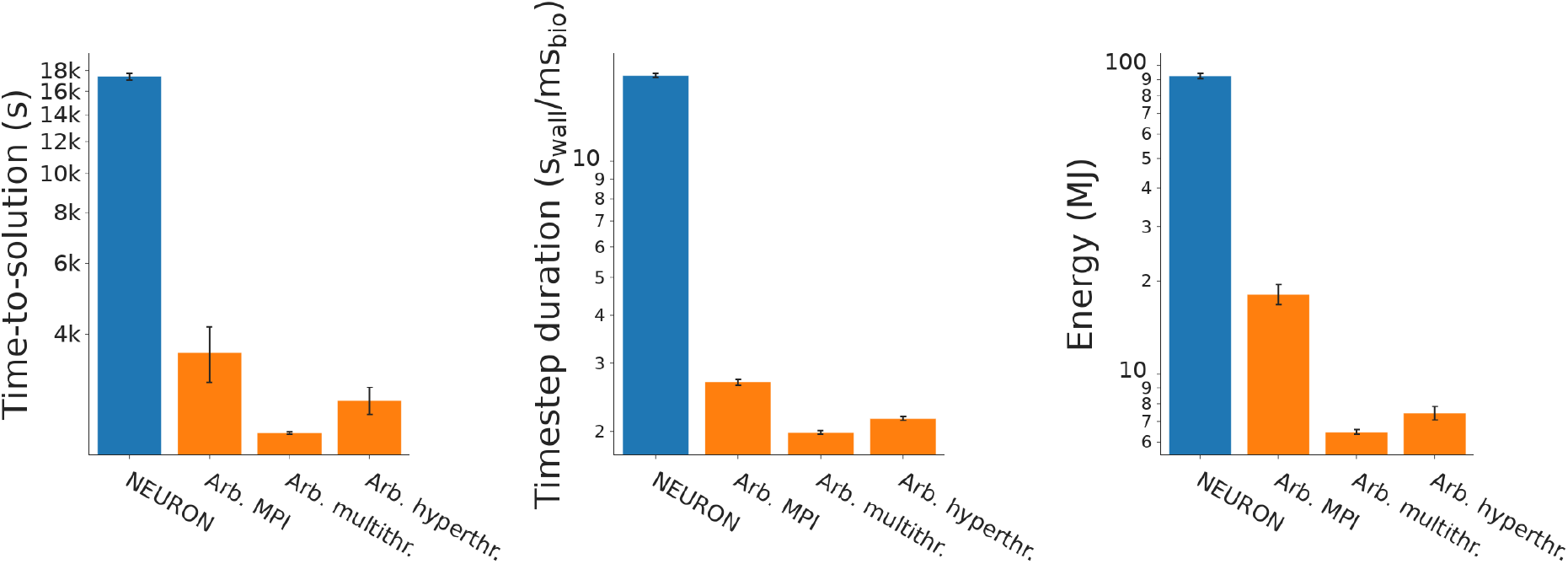
Distributed network model simulations. Time-to-solution (left), timestep duration (center), and energy consumption (right).

The benchmarks using GPU-acceleration show strong and weak scaling effects at multiple levels: there is a portion of the entire task that can not be parallelized, out of control of the simulator, visible in the increasing time-to-solution when more than 12 GPU nodes are used (Fig. 5, left and table 1, *Arbor GPU*). Additionally, a part of the simulation itself can not be parallelized, visible in a drop of the GPU occupation, increase in consumed node hours (Suppl Fig. 3) and stagnation of the timestep duration (Fig. 5, center) with more than 12 GPU nodes, for a model of this size. Thus, in terms of time-to-solution, 12 GPUs is the optimum size. If energy consumption is the main concern, using up to 4 GPU nodes is optimal (Fig. 5, right).

**Figure 5.**
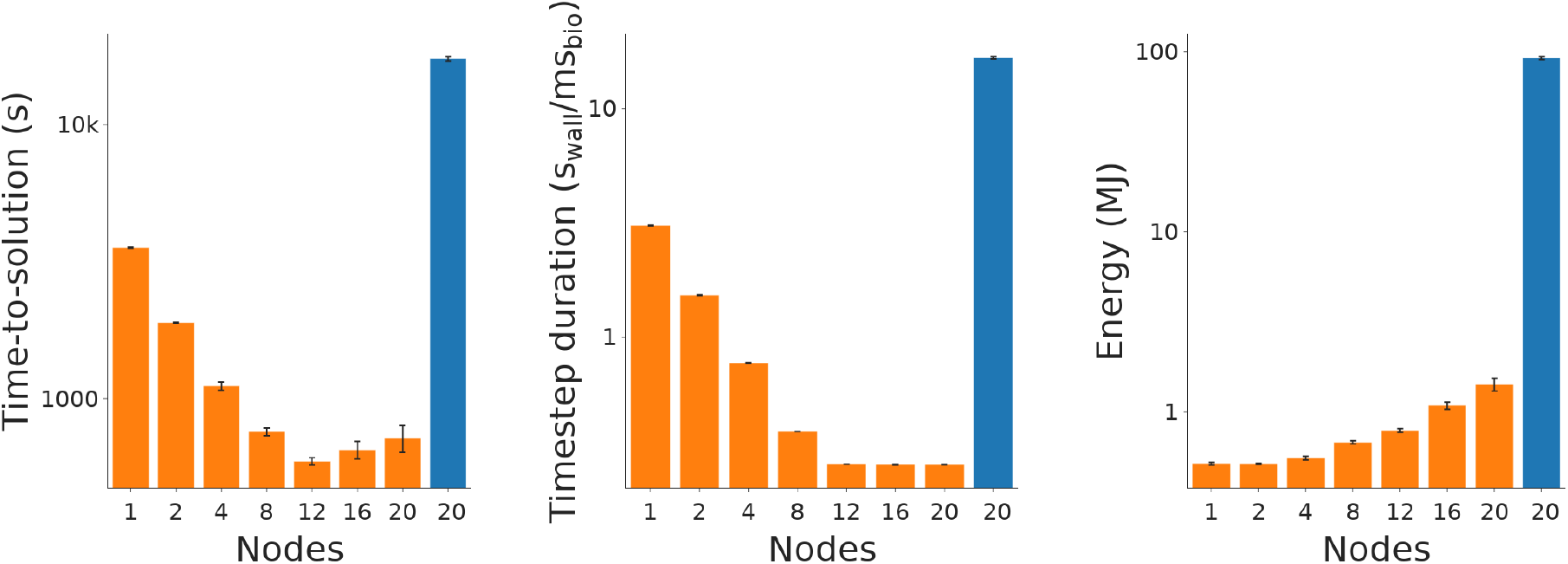
GPU-accelerated network model simulations. Time-to-solution (left), timestep duration (center), and energy consumption (right) in NEURON (blue, 36 CPU/node) compared to the GPU-accelerated network model simulations in Arbor (orange, 1 GPU/node). The NEURON simulation was ran only in the 20 CPU nodes (720 cores) configuration.

## 4 DISCUSSION &CONCLUSIONS

Even for *computational* neuroscientists, writing computer code is a means to an end. This has important implications for the quality of the code written, not just in terms of readability, style, performance, but also correctness, usability or shareability and longevity. In practice, scientific code is written once, and executed many times. Research groups tend to develop codebases that snowball: code is added for newly required features, but rarely removed or rewritten, or even studied and understood by new team members. These ad-hoc frameworks represent lock-in: it is not feasible to rewrite them or replace important underlying components. Even if efforts are made to apply development practices current in software engineering, it is rarely a core competency of scientific contributors and achieve only so much in attaining goals such as improving maintainability (the ease with which code can be understood and amended) and performance (hardware tomorrow may require a different way of distributing work than it does today). Concretely, in the course of this work NMODL files have been encountered that were passed down the generations in various labs and only their scientific output is validated: design, quality or performance are not an objective to analyze and tooling to feasibly tackle these problems are lacking.

While imperfect or suboptimal implementations are the nature of academic codes, moving to better specified and faster environments is worth pursuing. Well thought out specifications reduce the skill required in programming and errors in implementation subsequently, as well as allow for a better transferability of code between people and research groups, improving the usefulness of and scrutiny over such code. The ability to reuse code over sometimes decades is appealing, and with properly specified formats and tools, it is feasible. Naturally, researchers will use what gets the job done, so tools that interoperate through well specific formats such as NeuroML (Cannon et al., 2014) and the upcoming ModECI^9^ enable tool developers to write comprehensive test suites, as is common in software development, ensuring a consistency over time of results, interpretation and conformance; all ingredients required for reproducibility and long term sustainability of the field (Almog and Korngreen, 2016).

New projects, having the benefit of a clean break, are informed by the existence of pre-existing tools and their ecosystems, and are indebted to earlier efforts. One of the lessons that went into the design of Arbor is to avoid tying neuroscience to hardware architectures by optimizing for (or even being designed around) specific hardware features. Arbor features a compartmentalized design, which allows users in the present and future to write optimized implementations for combinations of (mathematical models of) cells and hardware (“Cell Groups”). This is how Arbor currently lets users transparently shift computation between GPUs and CPUs, and it enables users to execute their neuroscientific models unchanged on future hardware. Naturally, clean breaks come with the cost of having to re-implement features existing elsewhere, or do without. Model databases will not be, at least not without a conversion tool or layer, usable with such new tools. Although it is argued that reusability is in practice low (Almog and Korngreen, 2016), the vast wealth of data compatible with NEURON presents a challenge to any such break with the past. In this paper it is demonstrated that extending existing frameworks such as the BSB to be compatible with new tools or formats is feasible, and hopefully a small step towards evolving the ecosystem.

In conclusion, the comparison in this publication shows Arbor to be more efficient than NEURON under every condition and metric, up to 400 times in undistributed benchmarks, and 60 times in large scale distributed benchmarks. However, the validation shows that the expected biological outcome isn’t matched exactly in functional terms; it is likely that the translation of the NMODL files, interpretation of the morphology files, differences in discretization and in default, implicit or hidden parameters will lead to differences in outcome. To closely match the output of the cell and synapse models to biology, optimization would be required. Although model optimization is possible on simulation agnostic tools, Arbor stands to benefit from providing its own single cell optimization tools so that scientists can smoothly transition to a new simulation environment without changes to their model output.

## FUNDING

The research leading to these results has received funding from the European Union’s Horizon 2020 Framework Programme for Research and Innovation under the Specific Grant Agreements No. 945539 (Human Brain Project SGA3). R. DS is a recipient of a PhD fellowship from the University of Pavia.

## DATA AVAILABILITY STATEMENT

The sources, code and scripts used to produce the figures and data in this publication are freely available at https://github.com/Helveg/arb-nrn-comp.

---

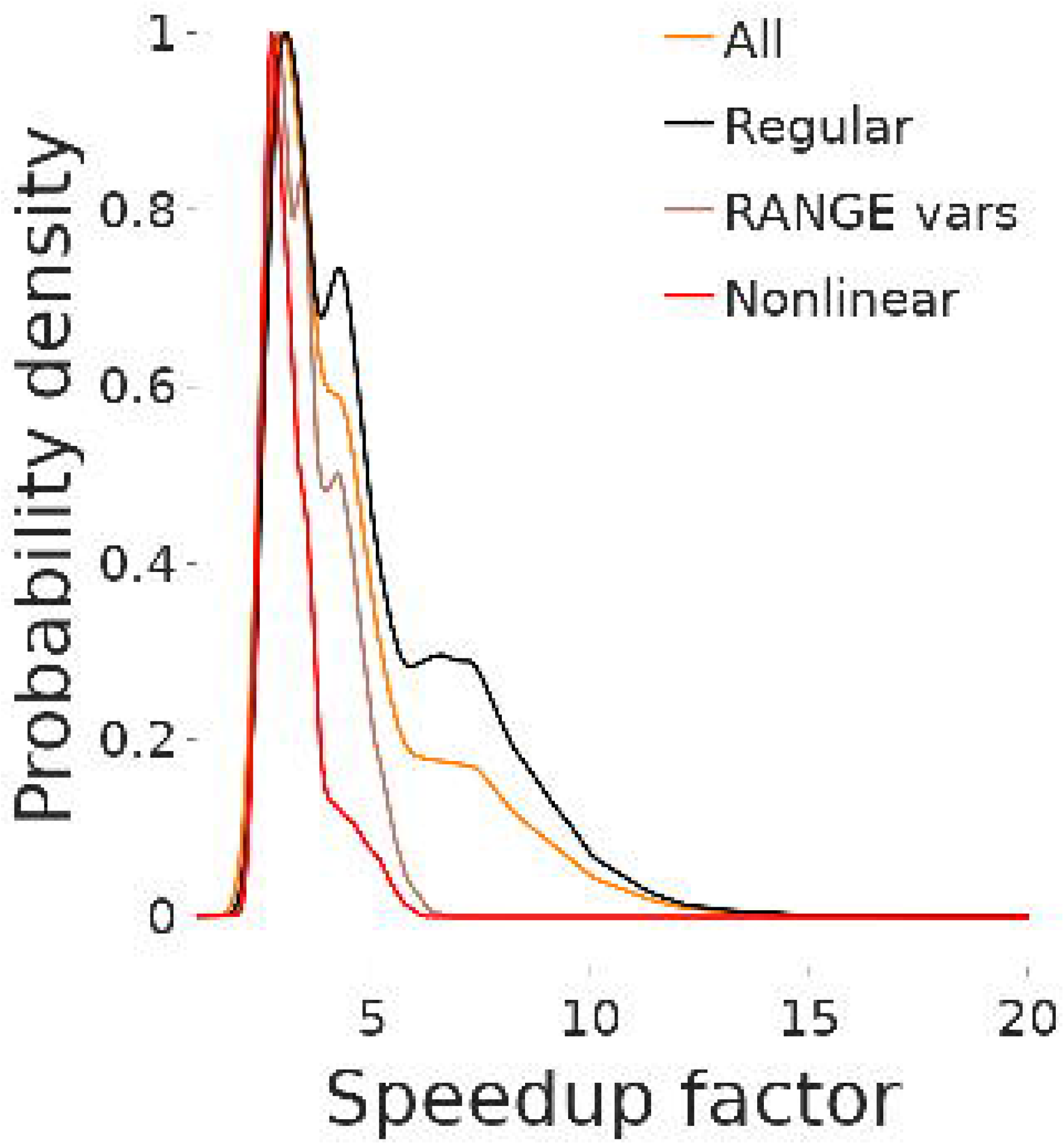

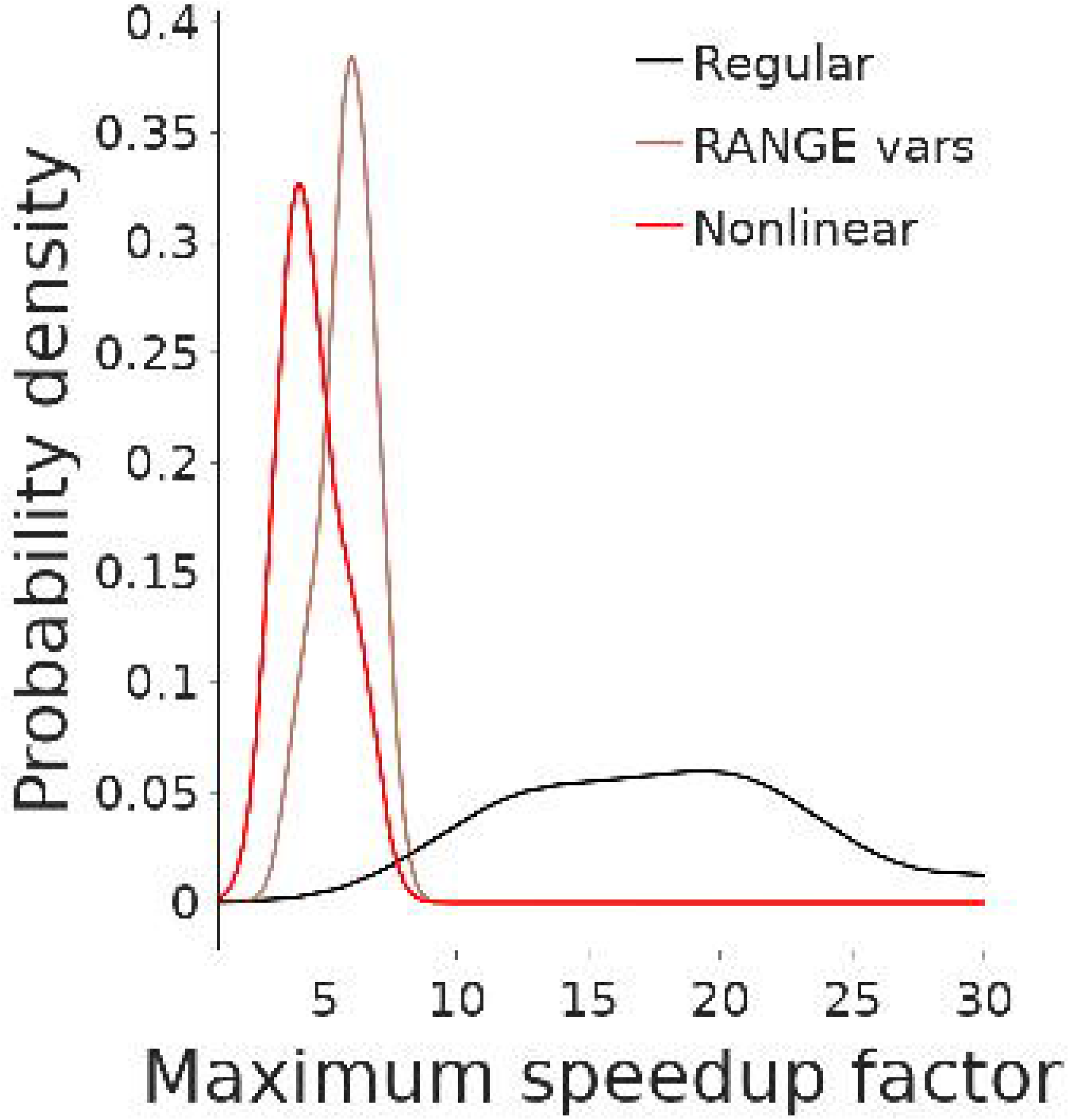

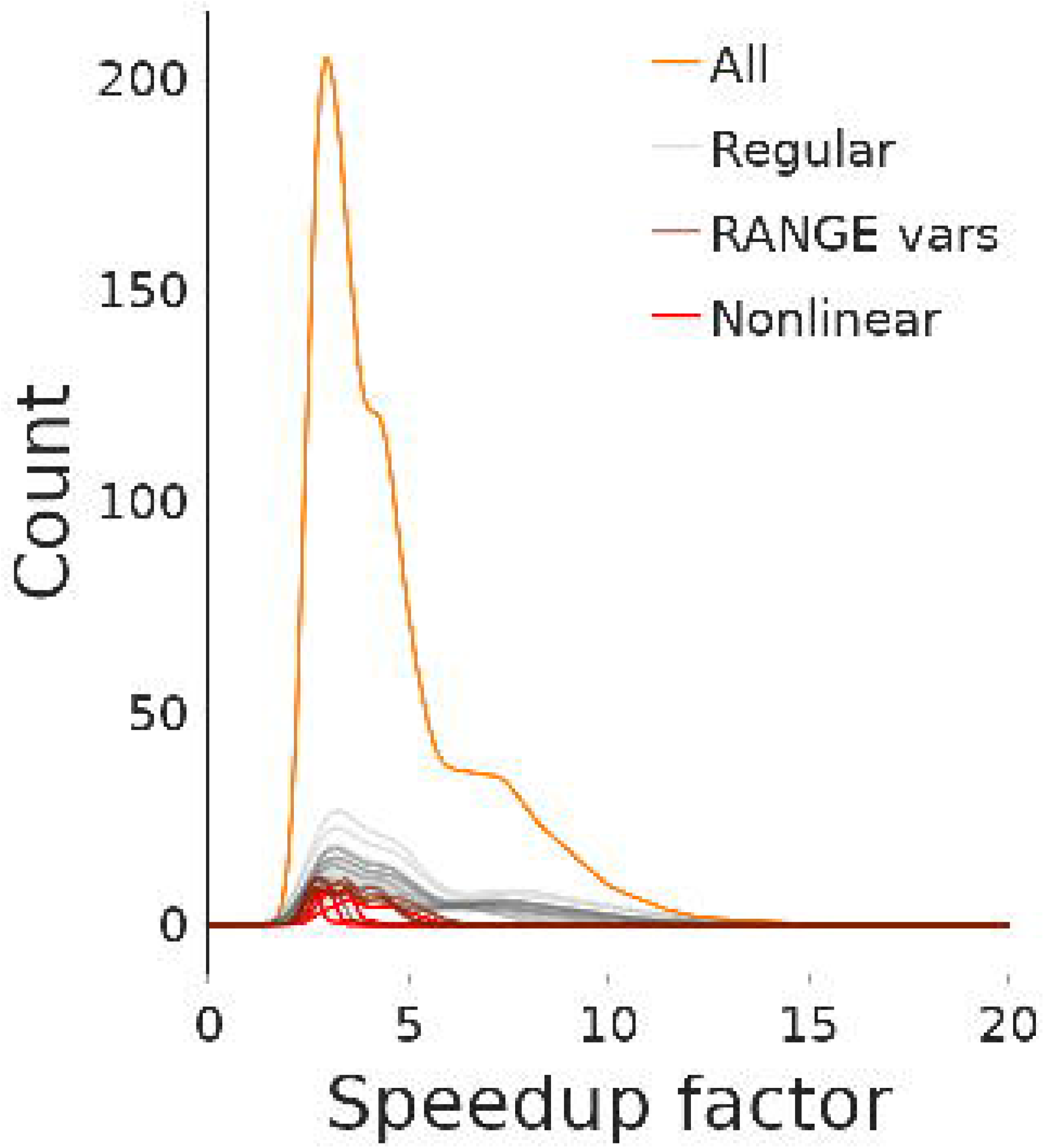

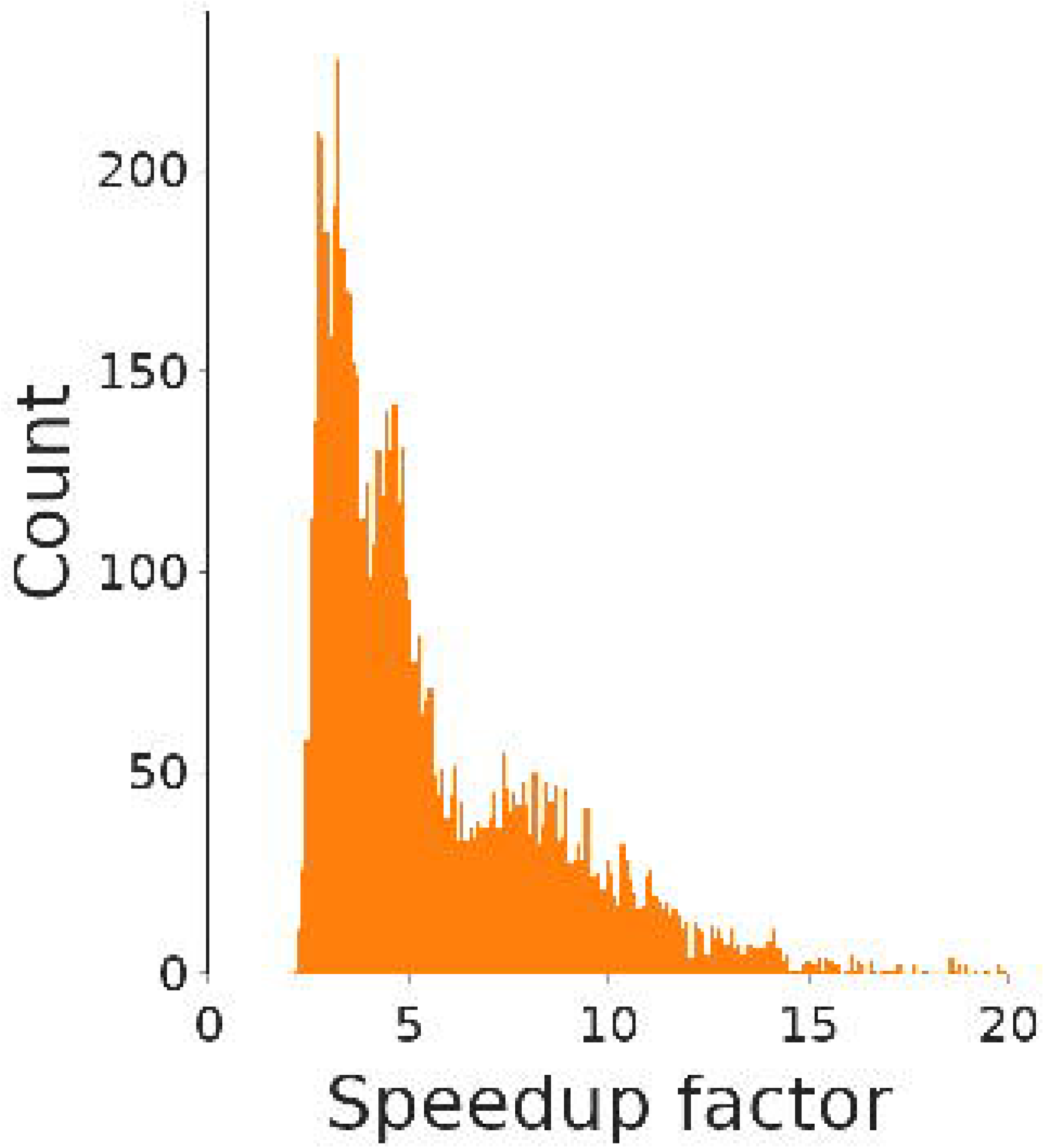

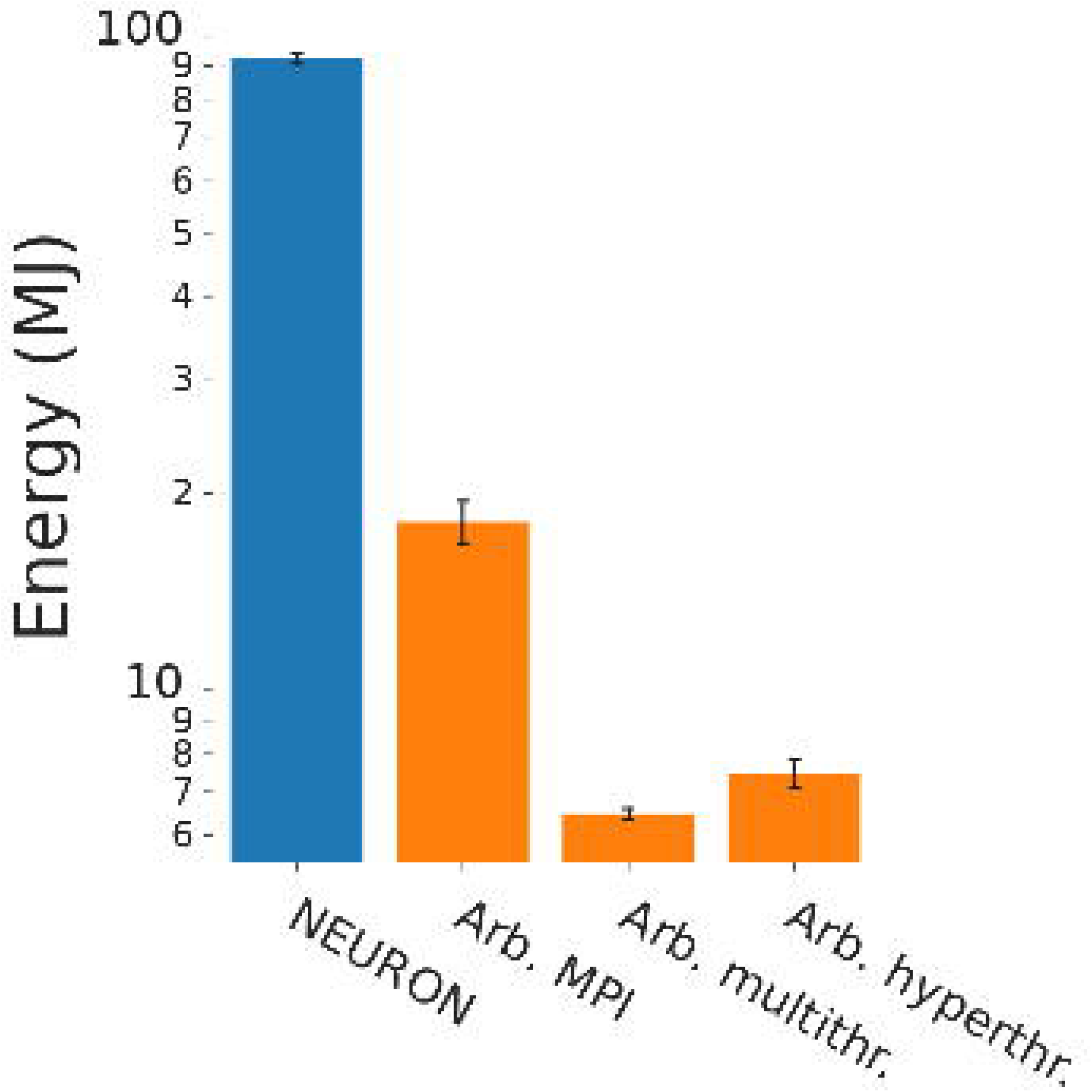

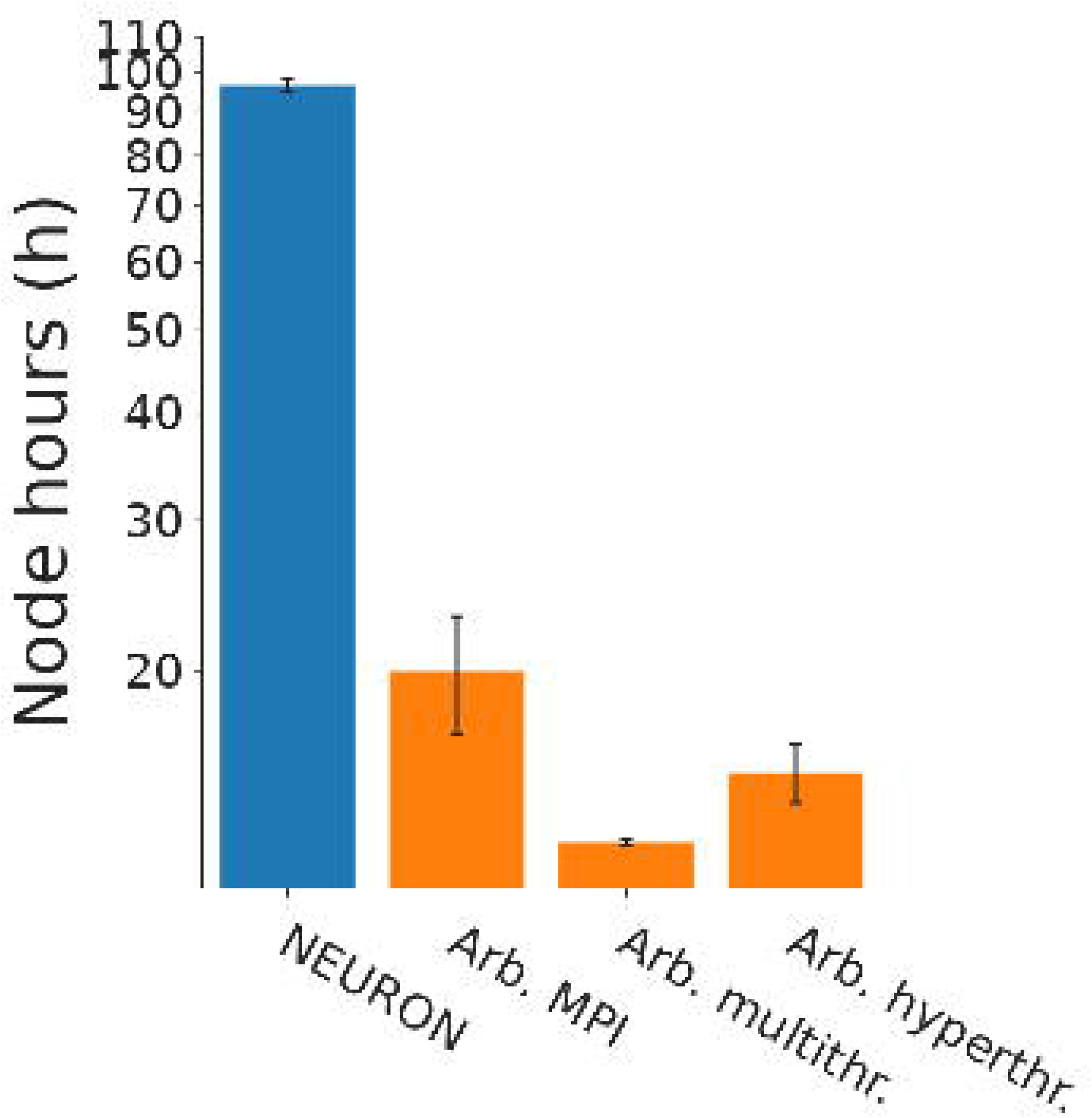

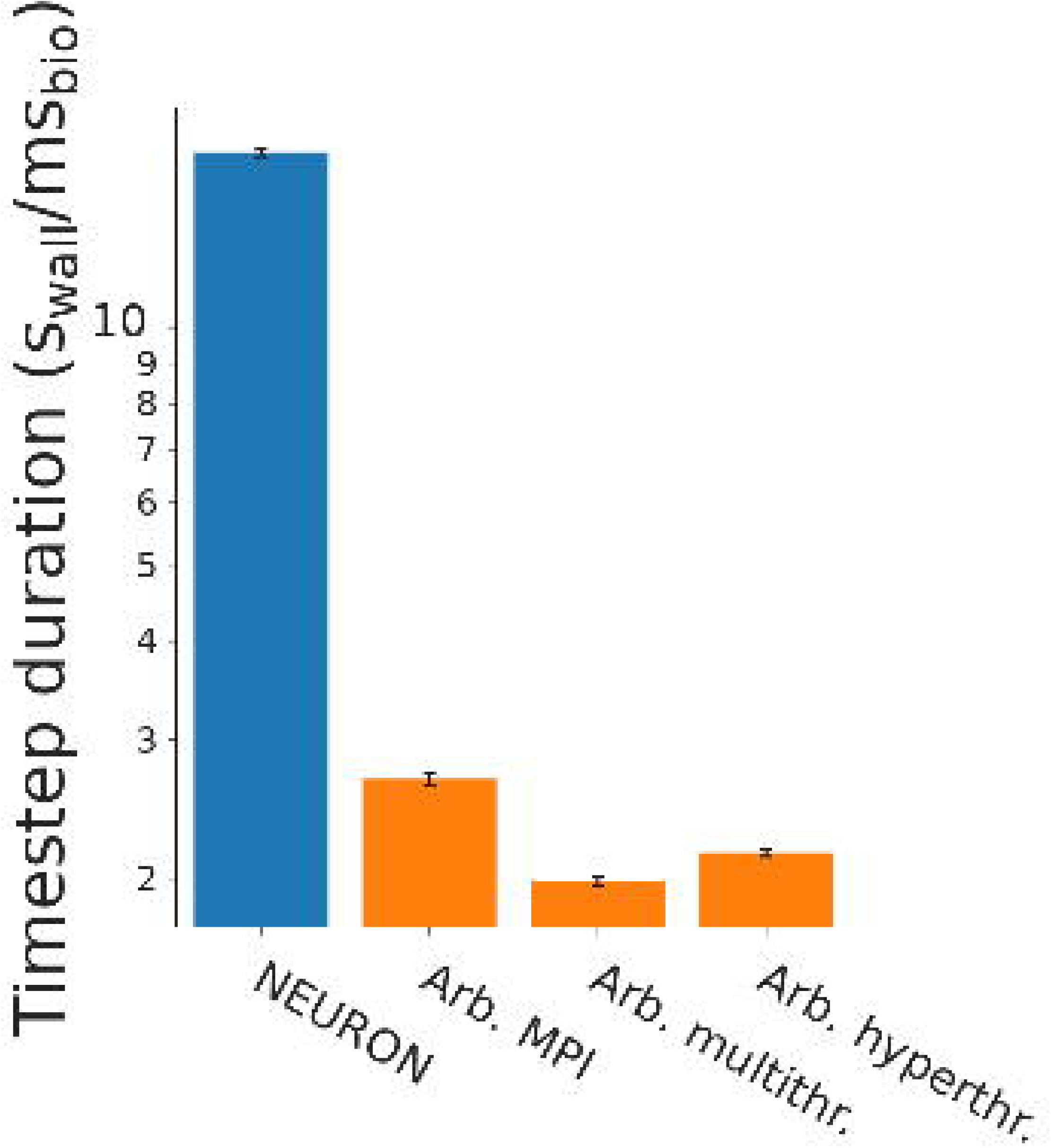

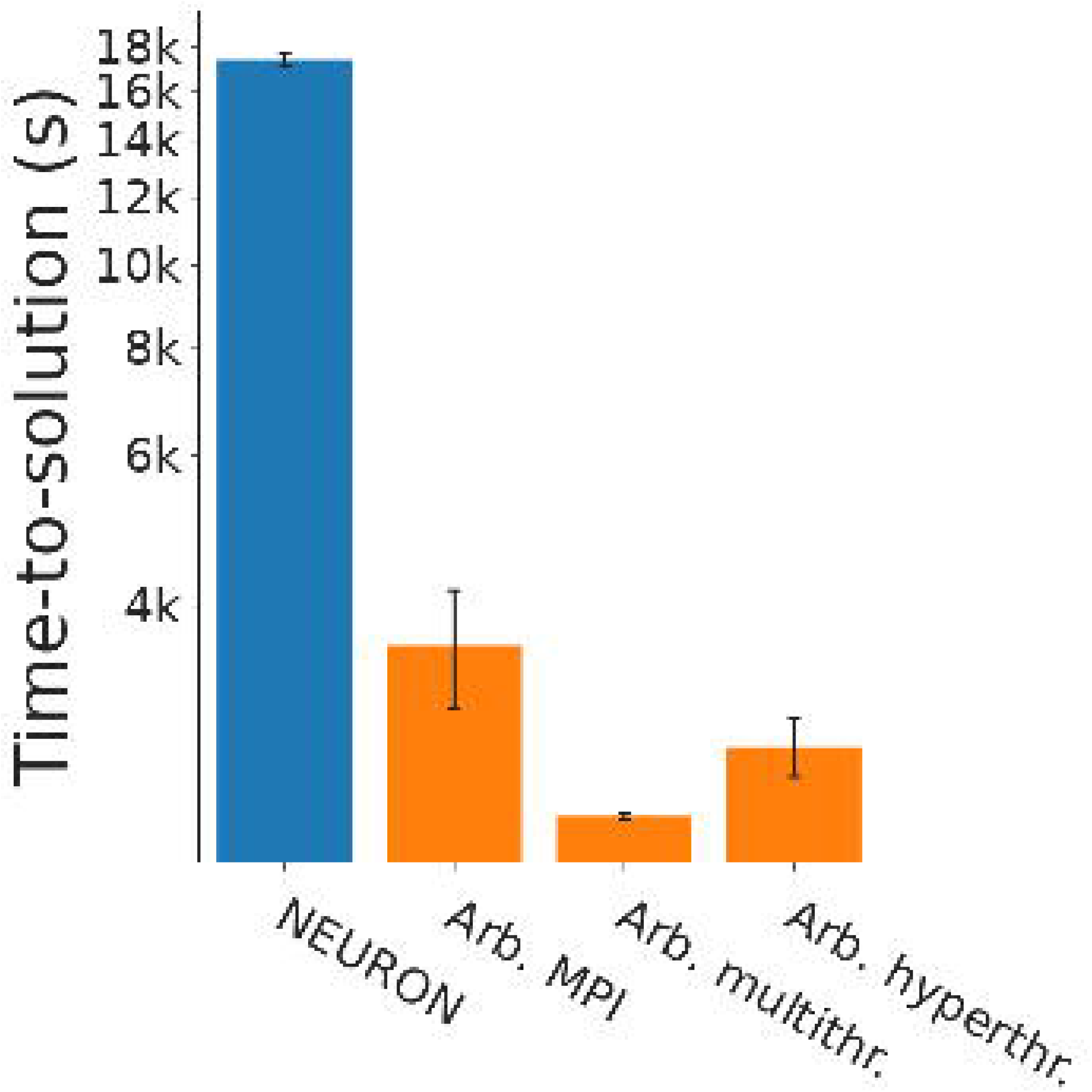

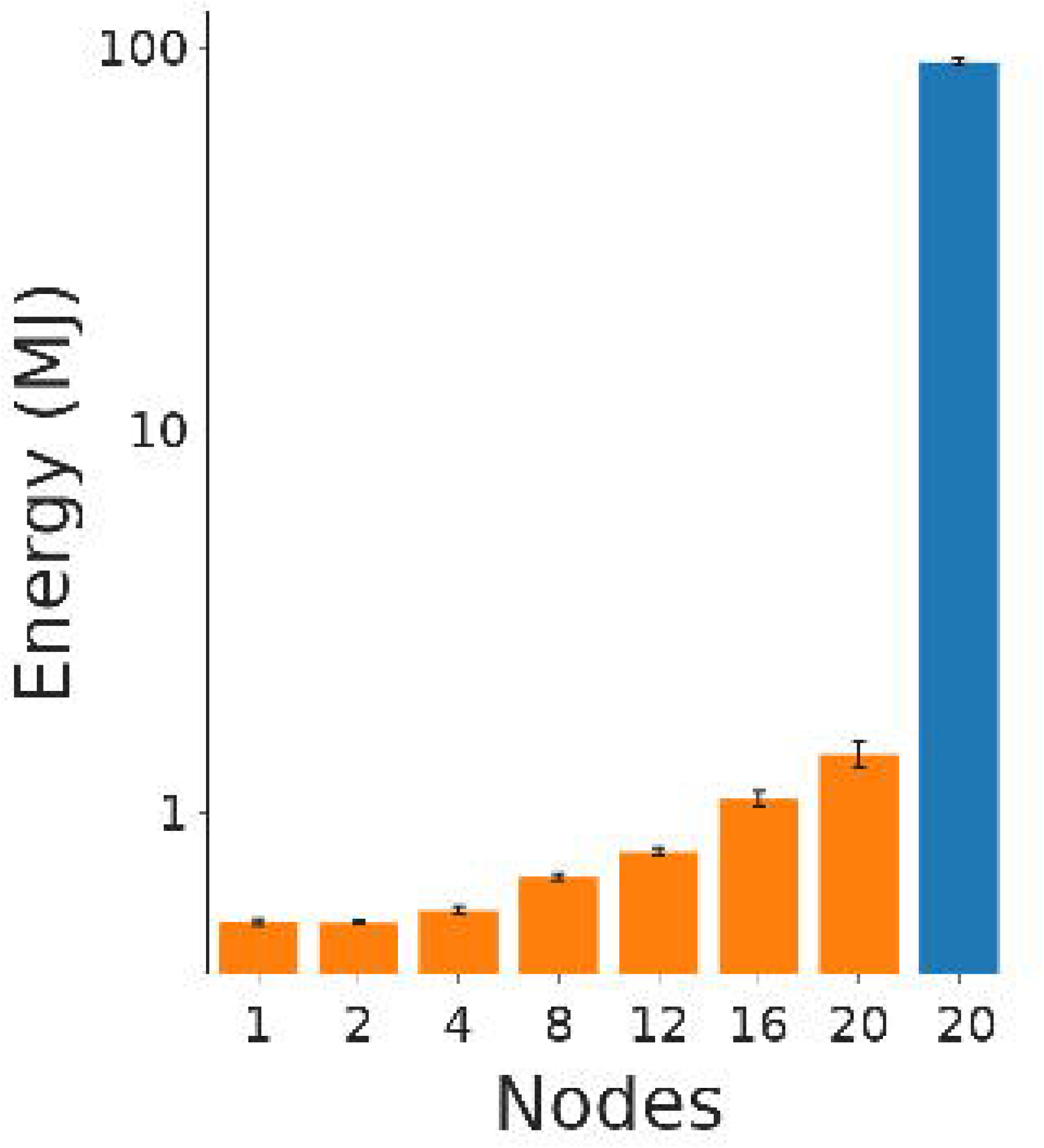

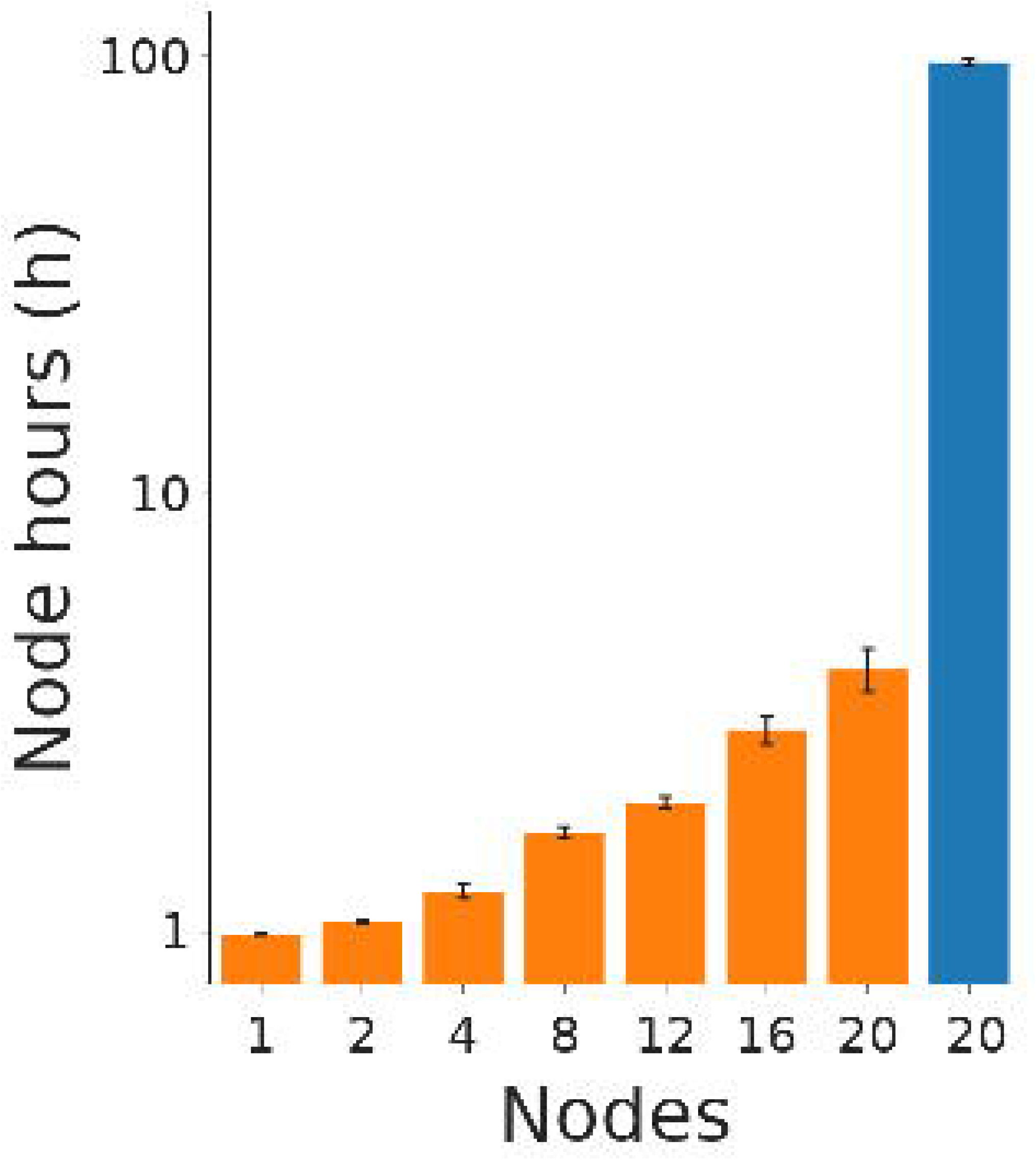

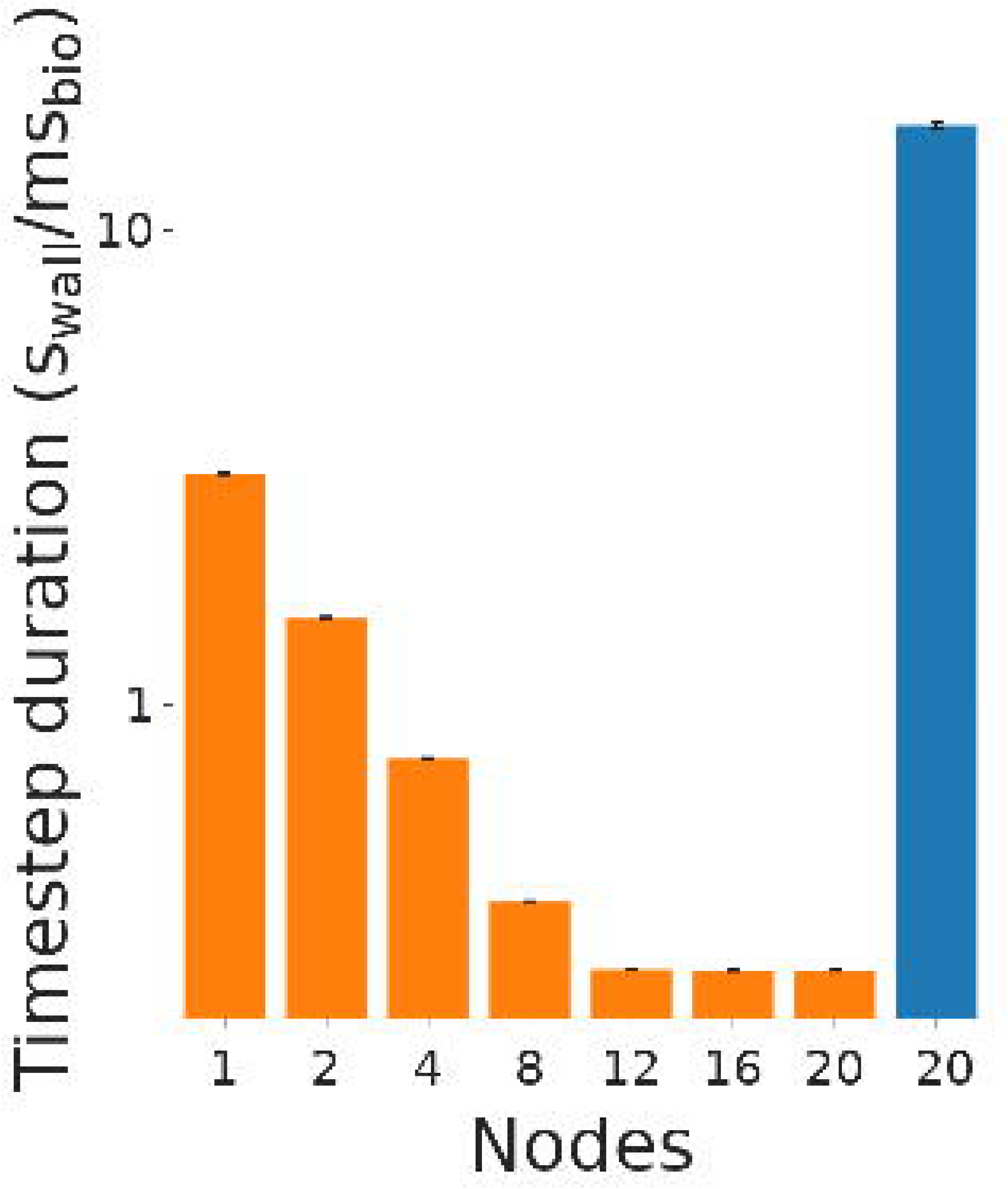

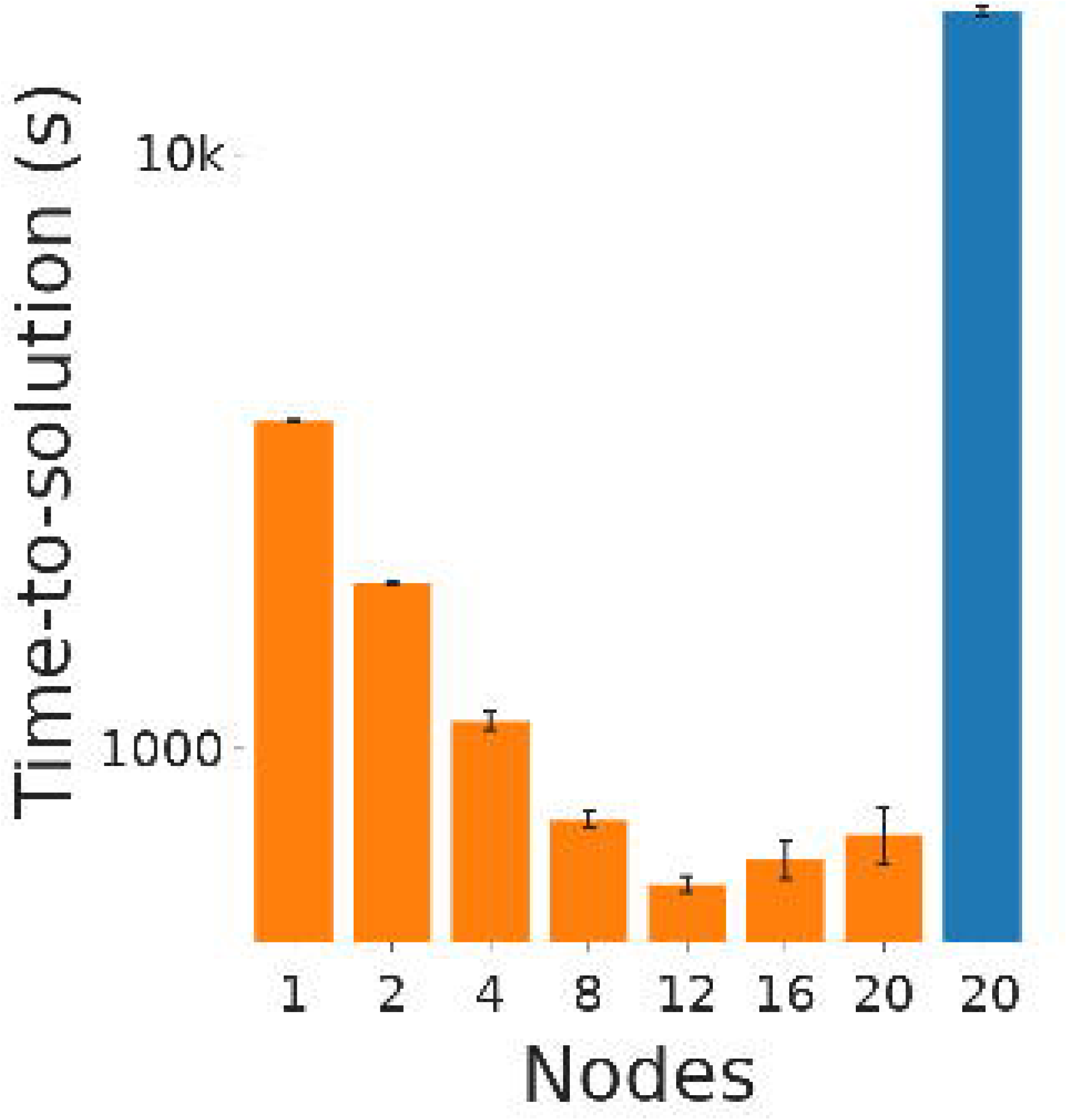

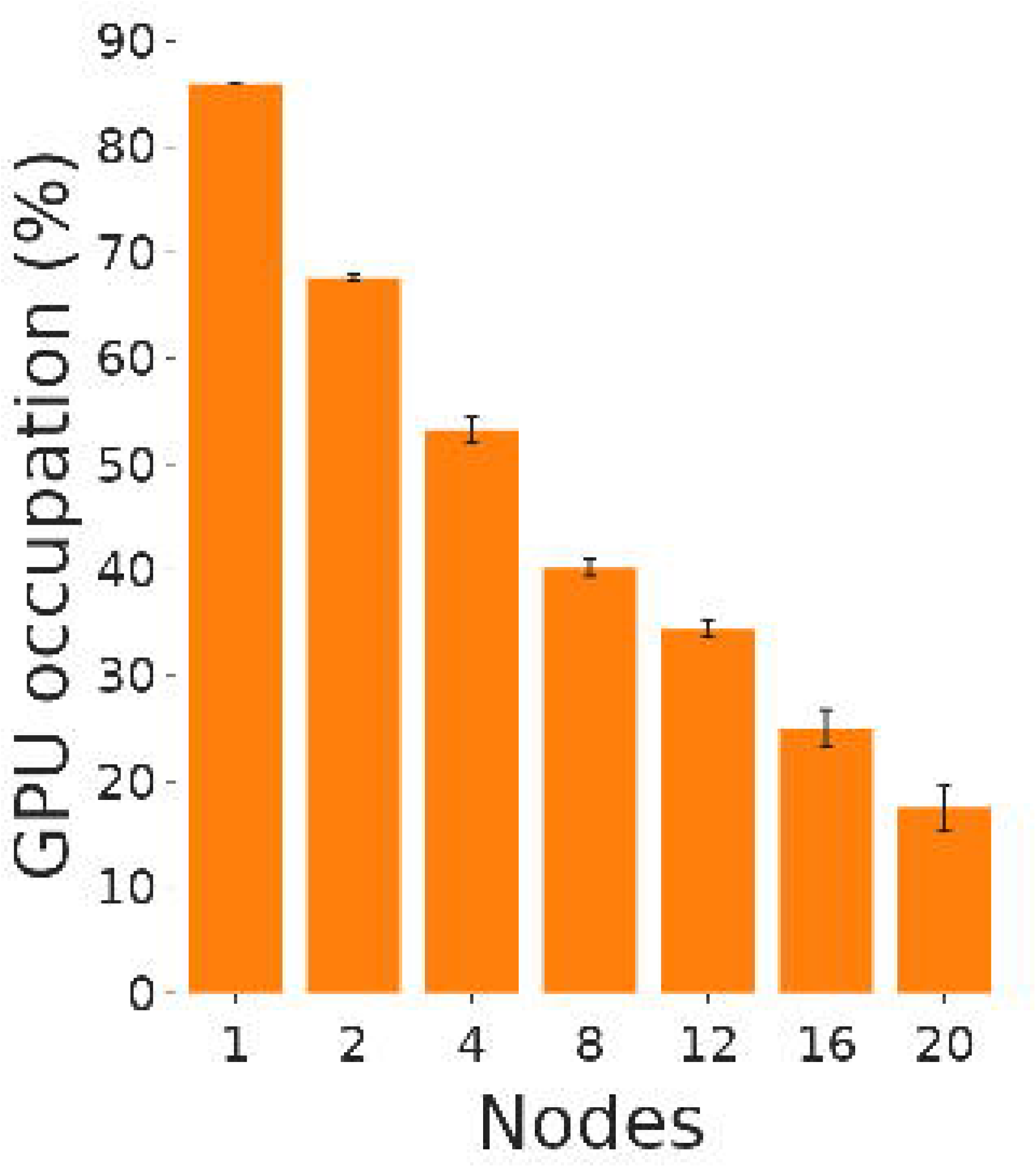

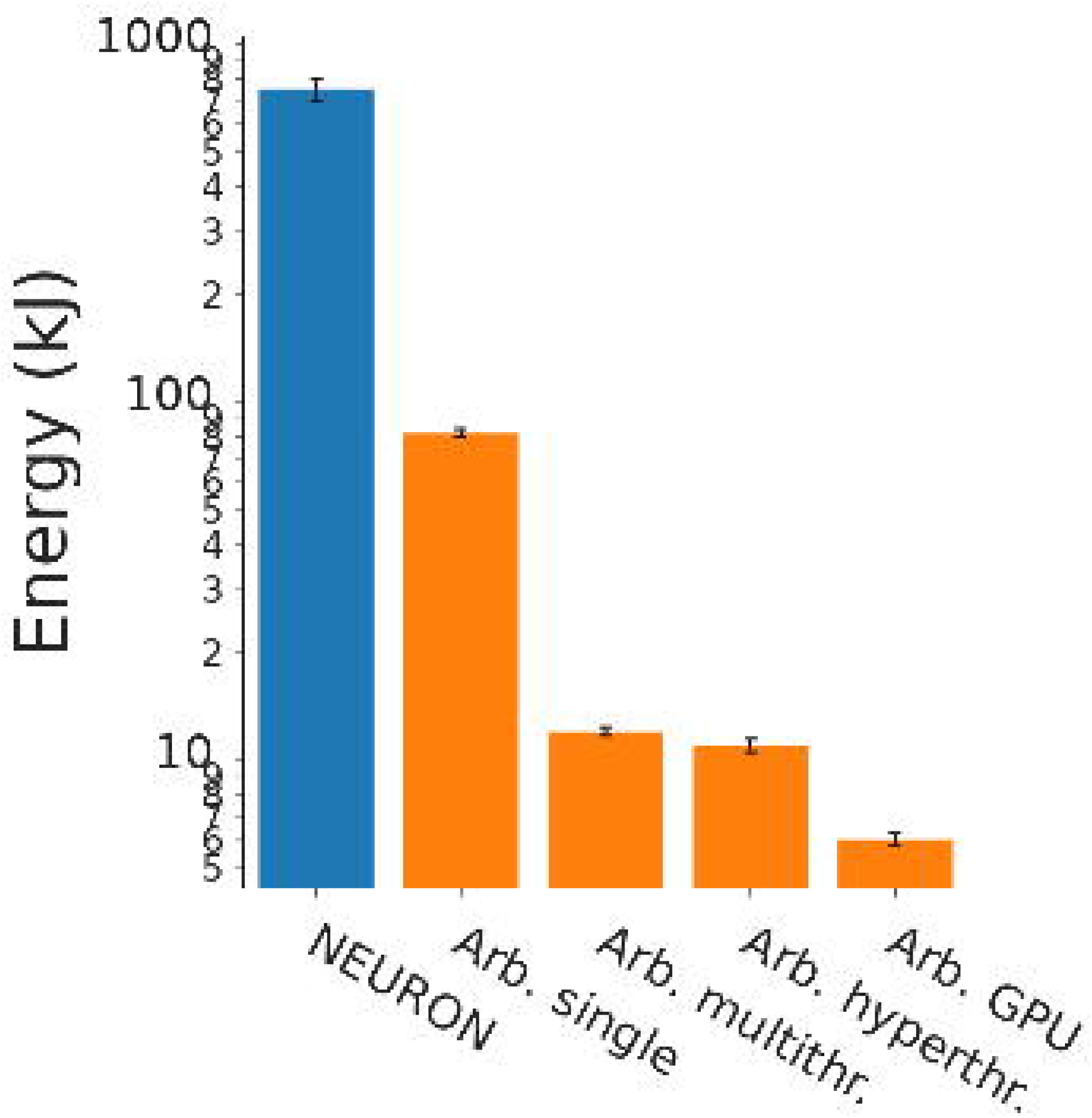

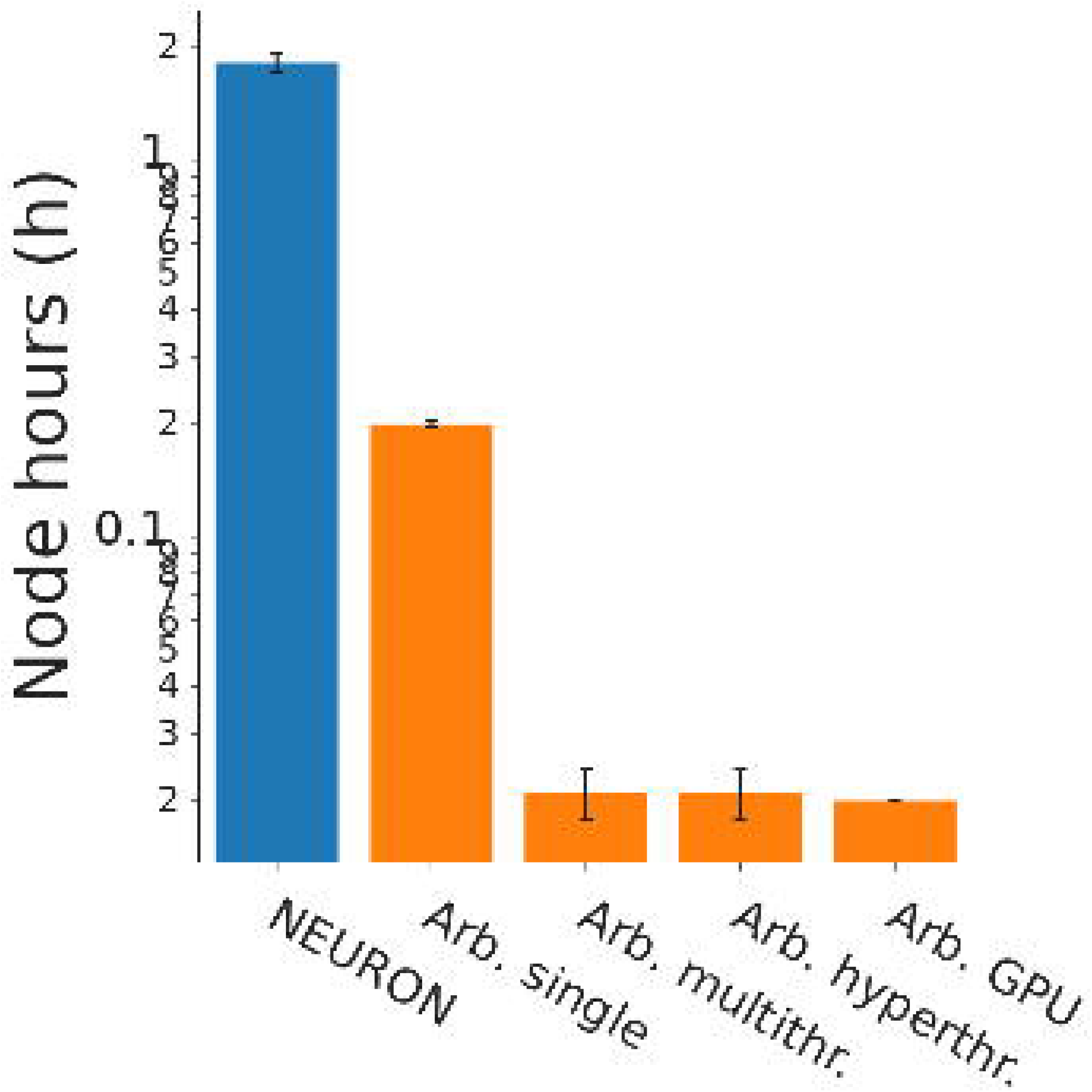

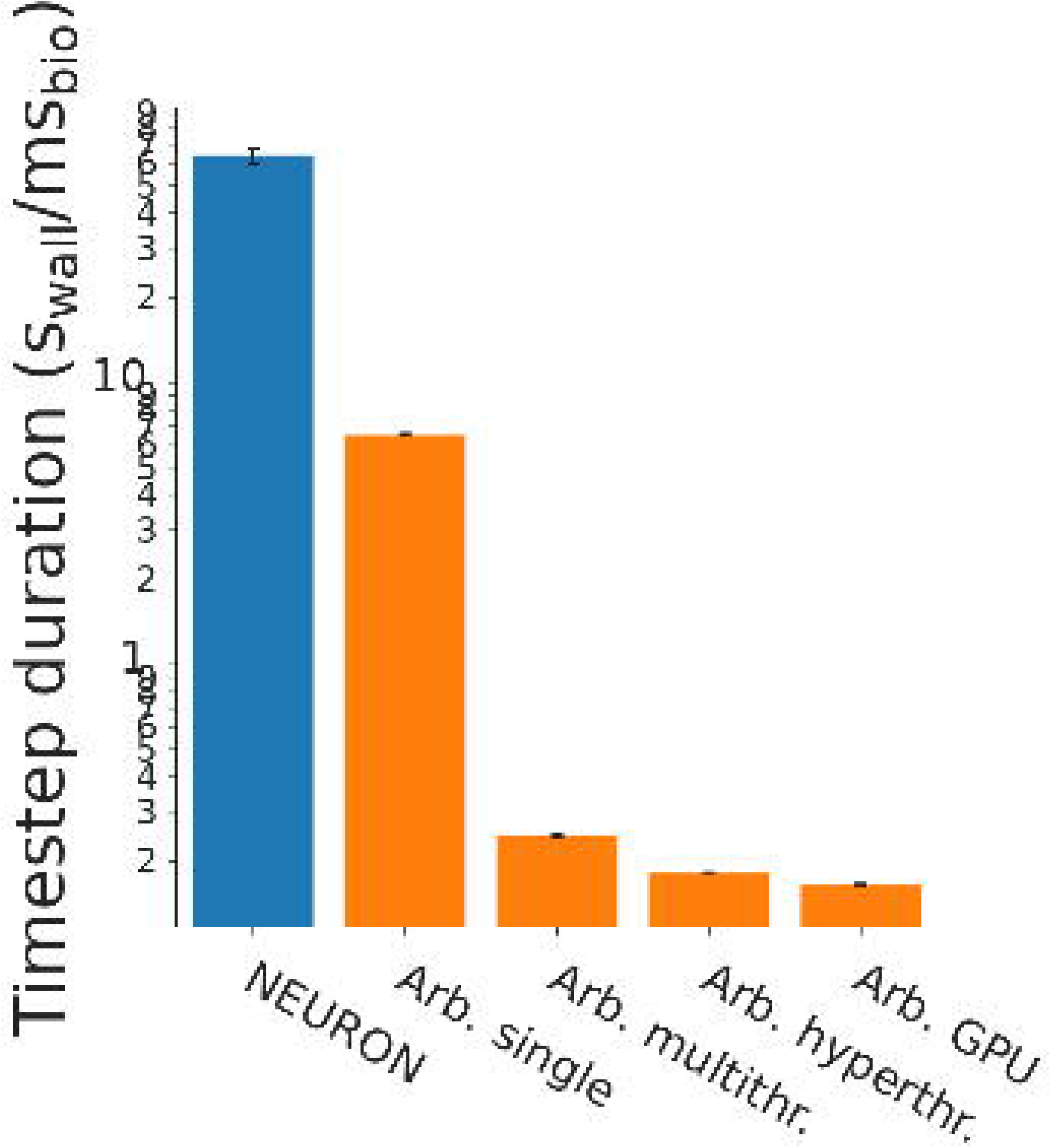

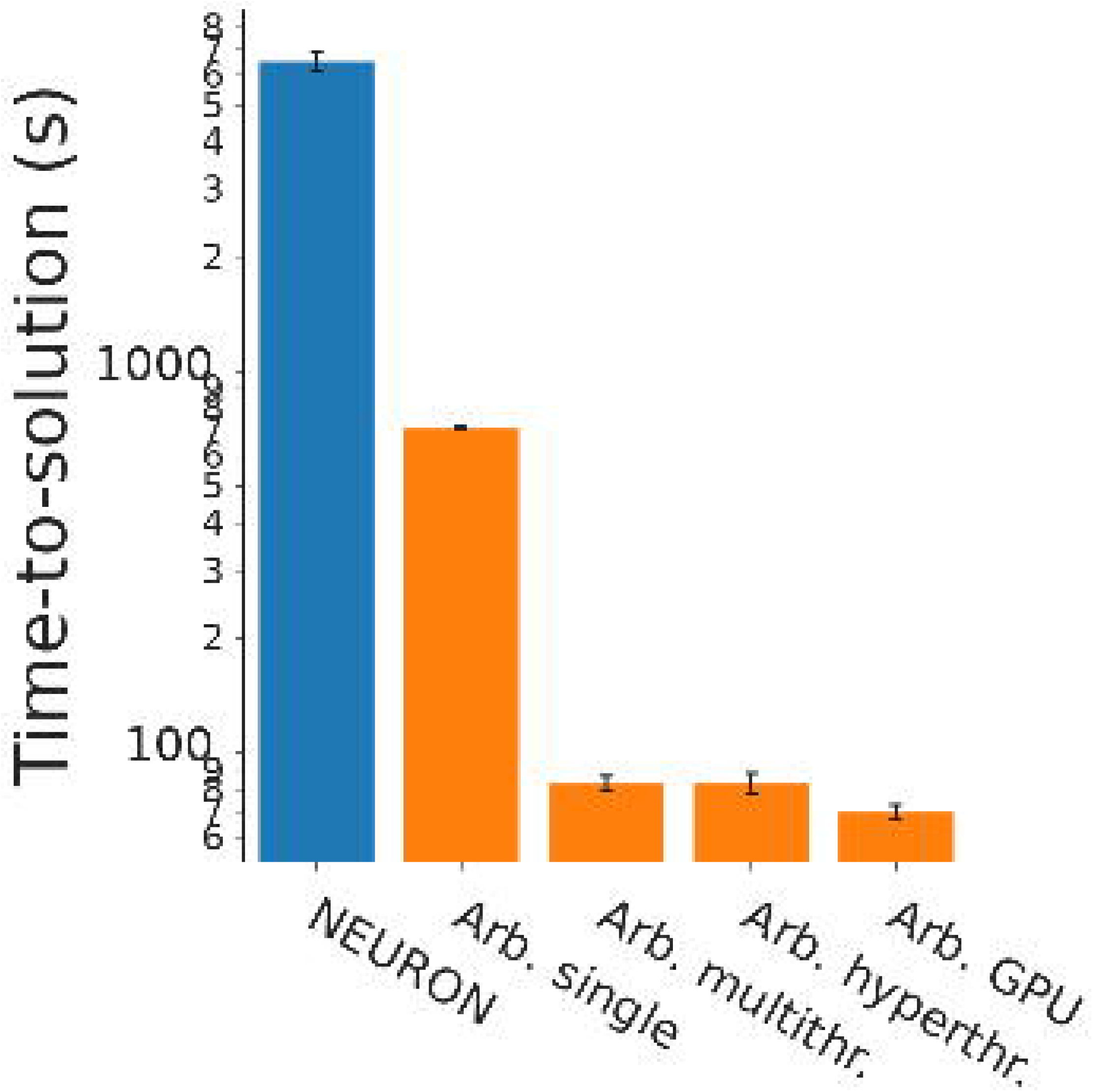

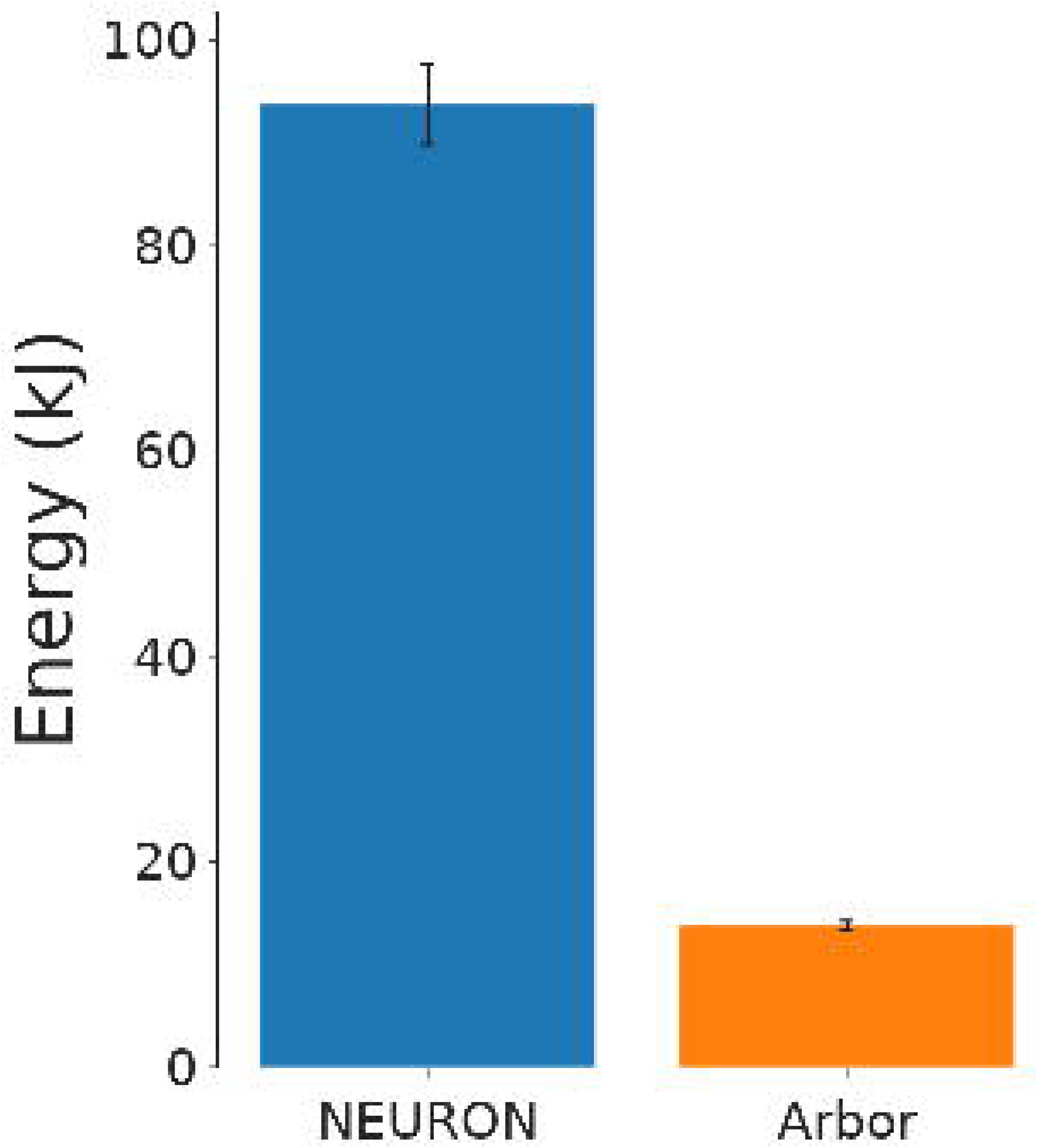

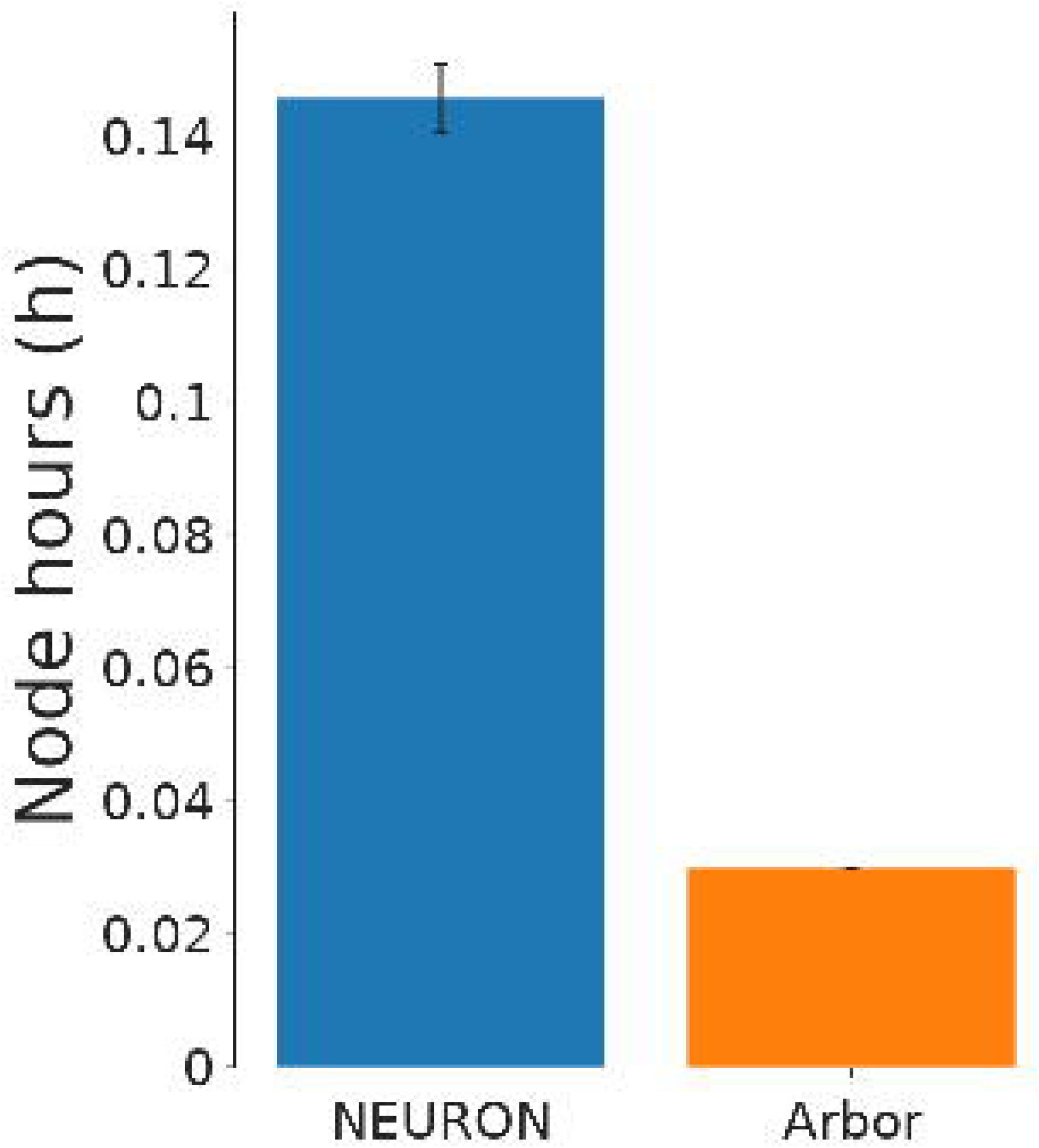

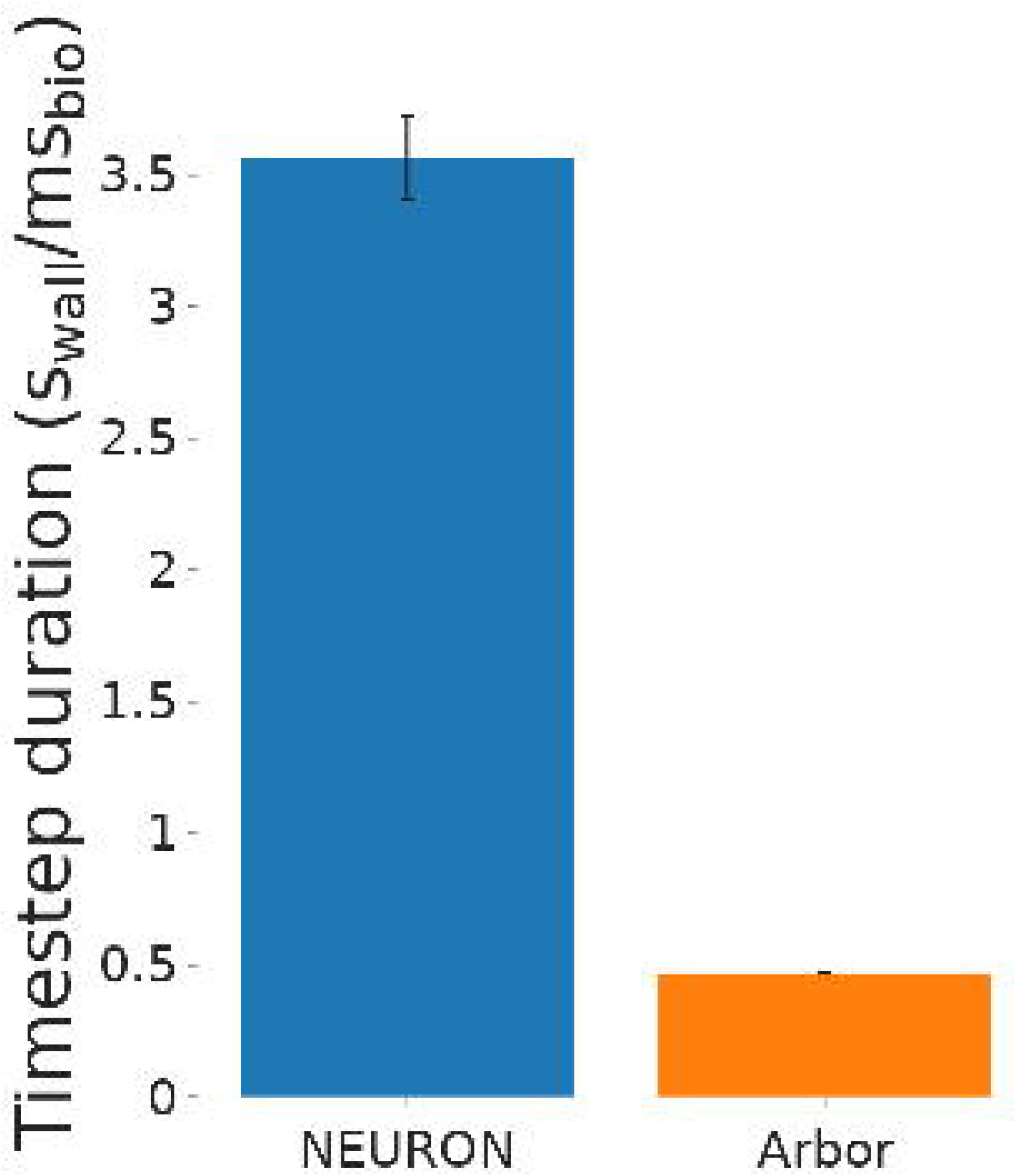

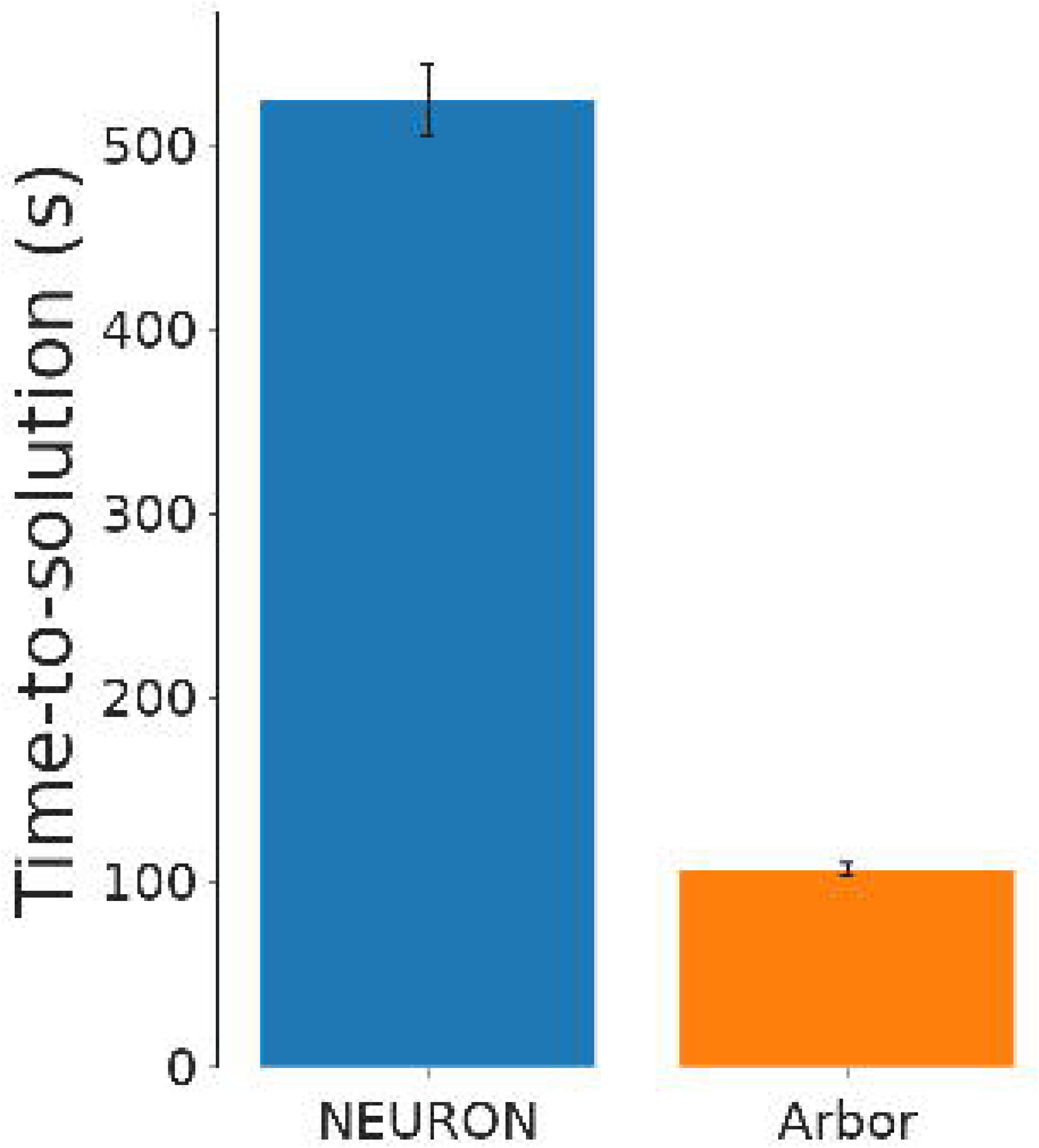

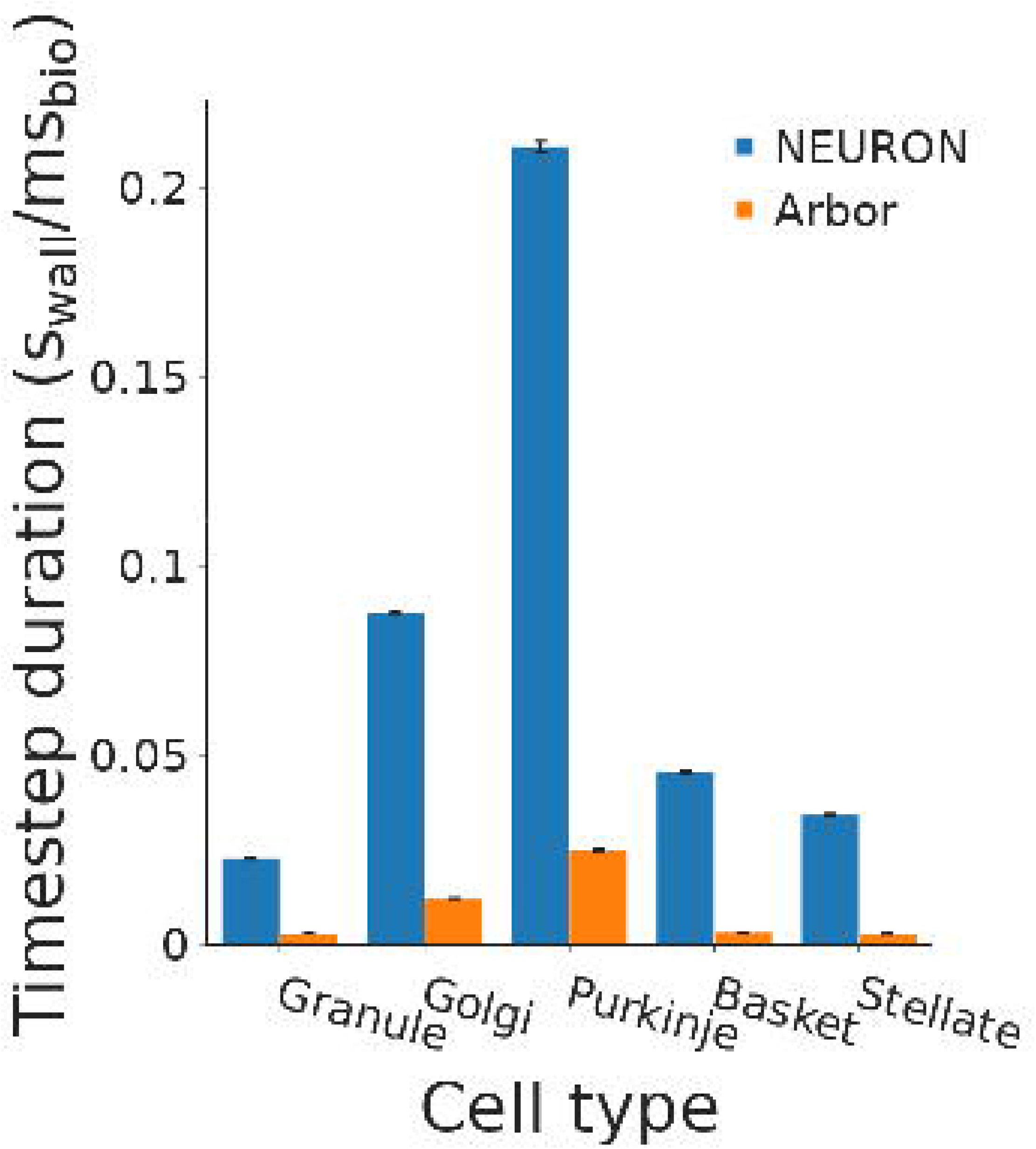

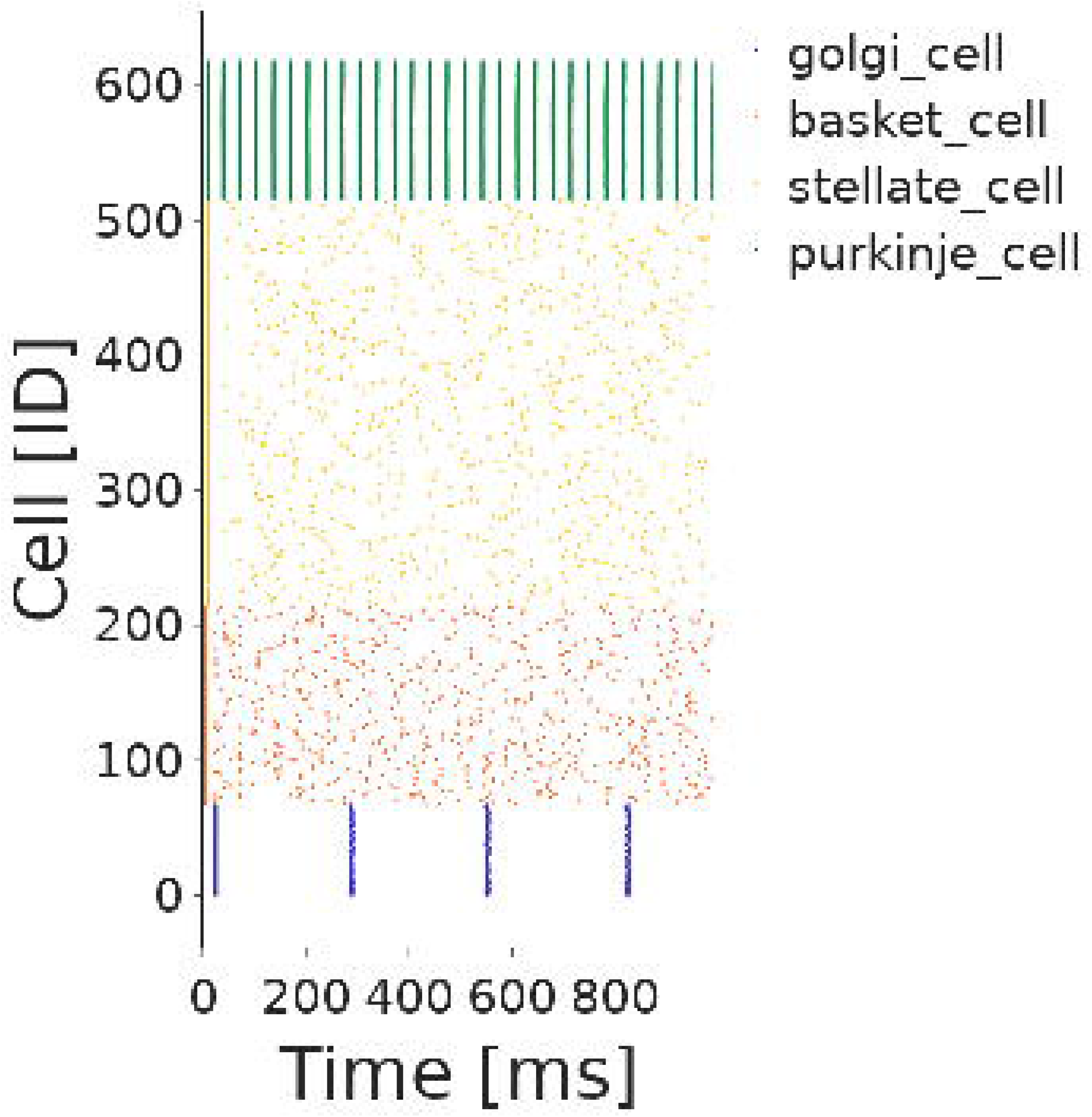

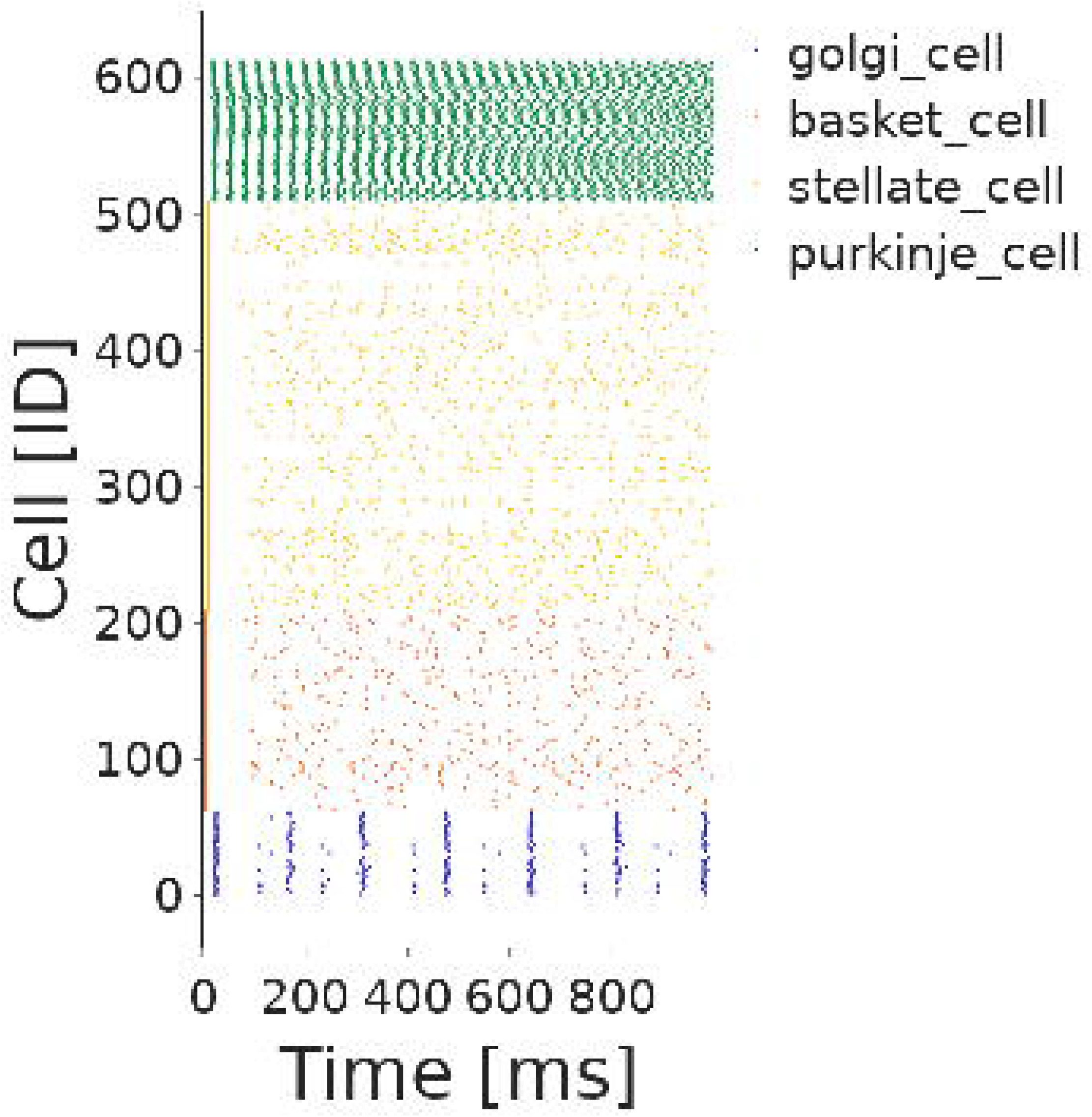

## Footnotes

1 github.com/dbbs-lab/dbbs-mod-collection

2 Arbor NMODL dialect guidelines

3 github.com/dbbs-lab/catalogue

4 github.com/dbbs-lab/models

5 The basket cell model is an unpublished adaptation of the stellate cell model

6 github.com/dbbs-lab/arborize

7 PizDaint is a supercomputer which is a hybrid Cray XC40/XC50 system and is the flagship system for the Swiss national HPC Service. For more details, see https://www.cscs.ch/computers/piz-daint/

8 Most of the RANGE variables were unused, or did not need to be RANGE variables and represent overheads incurred by *bitrot*, improper maintenance of source code over time.

9 modeci.org

## Notes

https://github.com/Helveg/arb-nrn-comp

### Competing Interest Statement

The authors have declared no competing interest.

