## Supplementary Material for "Cross-comparison of state of the art neuromorphological simulators on modern CPUs and GPUs using the Brain Scaffold Builder"

### 1 SUPPLEMENTARY TABLES AND FIGURES

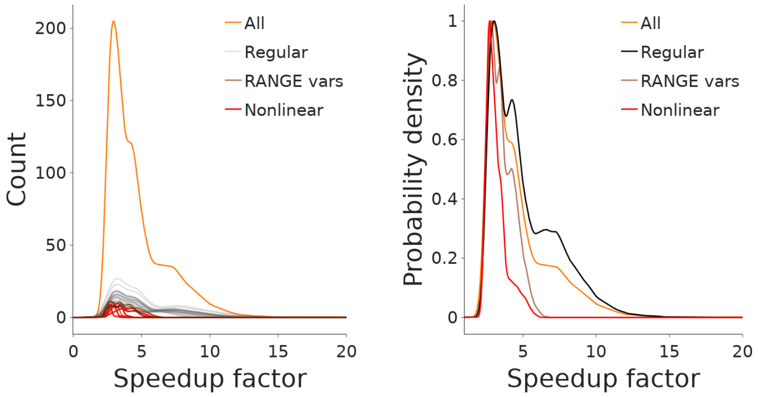


**Figure S1.** Left: Count (smoothed lines of histograms obtained through KDE) for the speedup factors of each of the 33 individual mechanisms when they occur in one of the 46,939 benchmarks; i.e. each line represents how a mechanism is likely to impact performance. Grouping based on a-posteriori inspection of the NMODL files. Right: Normalized probability density (through KDE) of each mechanism group, compared to the probability density of all mechanisms

| Model | Electrical nodes | Regular mech. | RANGE mech. | Nonlinear mech. | Est. weight |
|---|---|---|---|---|---|
| GranuleCell | 114 | 241 | 224 | 114 | 1245.6 |
| GolgiCell | 227 | 791 | 292 | 227 | 2406.3 |
| PurkinjeCell | 467 | 3815 | 462 | 462 | 6771.8 |
| BasketCell | 114 | 460 | 72 | 114 | 1084.6 |
| StellateCell | 120 | 619 | 31 | 120 | 1164.5 |

**Table S1.** Analysis of the single cell models in terms of computational load. The electrical nodes (*segments* in NEURON and *control volumes* in Arbor) represent the cell model size and are the number of points in which the system of electrical equations have to be solved. The mechanism columns indicate the total number of mechanisms that are spread across those nodes. The estimated weight multiplies the number of mechanisms of each category with a weight (1, 2.5 and 3.9 respectively) obtained from the mechanism speedup factors and sums them together to represent model complexity.

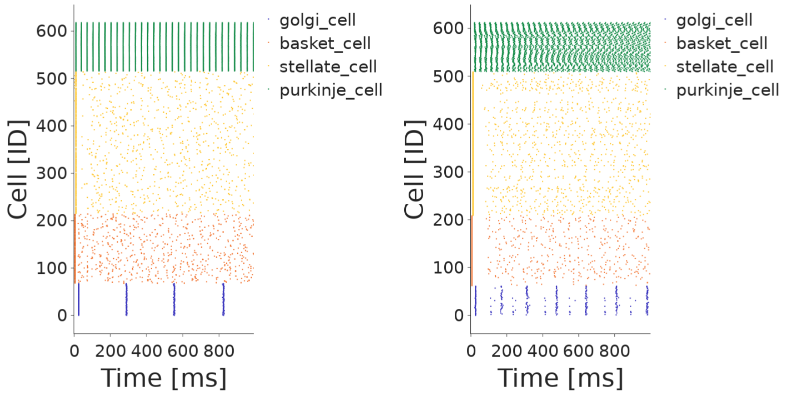


**Figure S2.** Left: Raster plot of an Arbor simulation. Right: Raster plot of a NEURON simulation. There are some differences in the spread of the spike times of the Golgi and Purkinje cell, but since the average firing rates were mostly the same this wasn't considered an issue for comparisons of the computational load of the benchmarks. Average firing frequencies are (reported as Arbor/NEURON values): 0.0/0.0 granule, 30.0/31.6Hz Purkinje, 4.0/4.85hz Golgi, 7.7/3.9Hz basket, 4.29/4.39 stellate. The quiescent granule cells are not shown.

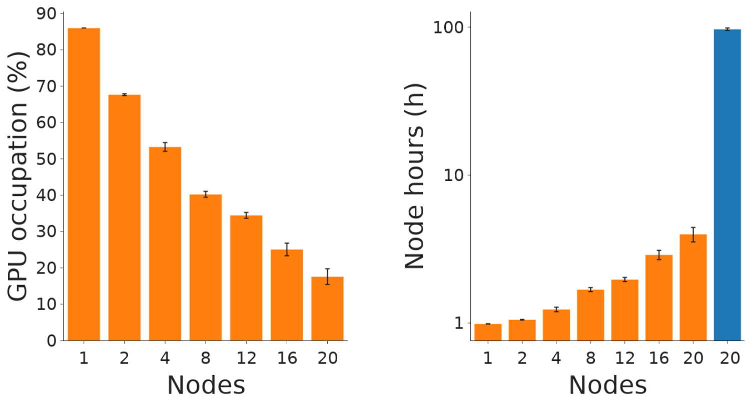


**Figure S3.** GPU accelerated results. Left: Average load for each GPU used in the simulation. Right: Total node hours used by the simulation. NEURON simulations used 20 CPU nodes (720 cores)
